# Combining pairwise structural similarity and deep learning interface contact prediction to estimate protein complex model accuracy in CASP15

**DOI:** 10.1101/2023.03.08.531814

**Authors:** Raj S. Roy, Jian Liu, Nabin Giri, Zhiye Guo, Jianlin Cheng

## Abstract

Estimating the accuracy of quaternary structural models of protein complexes and assemblies (EMA) is important for predicting quaternary structures and applying them to studying protein function and interaction. The pairwise similarity between structural models is proven useful for estimating the quality of protein *tertiary* structural models, but it has been rarely applied to predicting the quality of *quaternary* structural models. Moreover, the pairwise similarity approach often fails when many structural models are of low quality and similar to each other. To address the gap, we developed a hybrid method (MULTICOM_qa) combining a pairwise similarity score (PSS) and an interface contact probability score (ICPS) based on the deep learning inter-chain contact prediction for estimating protein complex model accuracy. It blindly participated in the 15th Critical Assessment of Techniques for Protein Structure Prediction (CASP15) in 2022 and ranked first out of 24 predictors in estimating the global accuracy of assembly models. The average per-target correlation coefficient between the model quality scores predicted by MULTICOM_qa and the true quality scores of the models of CASP15 assembly targets is 0.66. The average per-target ranking loss in using the predicted quality scores to rank the models is 0.14. It was able to select good models for most targets. Moreover, several key factors (i.e., target difficulty, model sampling difficulty, skewness of model quality, and similarity between good/bad models) for EMA are identified and analayzed. The results demonstrate that combining the multi-model method (PSS) with the complementary single-model method (ICPS) is a promising approach to EMA. The source code of MULTICOM_qa is available at https://github.com/BioinfoMachineLearning/MULTICOM_qa.

## 1. Introduction

Most proteins must fold into some specific 3D shape called tertiary structures to carry out their biological function. Two or more proteins (more precisely protein chains) often interact to form a protein complex or assembly to collaboratively perform a complicated biological function such as transduction of an extracellular signal into a cell to trigger gene expression. Therefore, predicting the tertiary structure of a single protein (monomer) and the quaternary structure of a protein complex/assembly (multimer) from its sequence has been a major pursuit of the scientific community for the last few decades because the experimental methods of determining protein structures are expensive and time-consuming and therefore can only be applied to a small fraction of proteins.

To rigorously and objectively assess the progress in protein structure prediction, the Critical Assessment of Techniques for Protein Structure Prediction (CASP) was launched in 1994 and continued to be held every two years (Moult et al., 1995; Kryshtafovych et al., 2014; Moult et al., 2016; Kryshtafovych et al., 2019; Kwon et al., 2021), with the latest CASP15 experiment concluded in 2022. Stimulated by the CASP experiments, significant progress has been made in protein tertiary structure prediction. Particularly, with the development of deep learning-based methods over the last decade (Eickholt & Cheng, 2012; Wang et al., 2017; Kandathil et al., 2019; Hou et al., 2019; Senior et al., 2019; Zheng et al., 2019; Senior et al., 2020; Yang et al., 2020; Baek et al., 2021;Jumper et al., 2021,Liu et al., 2022), the prediction of protein tertiary structures had steadily improved from one CASP experiment to another until AlphaFold2 (Jumper et al., 2021) demonstrated its outstanding capability of predicting tertiary structures with the quality close to that of the experimental techniques such as X-ray crystallography. The success of AlphaFold2 and other deep learning methods has revolutionised the prediction and application of protein tertiary structures in various biological research.

As the tertiary structure prediction problem has been largely solved, the field started to focus more on predicting the *quaternary* structures of protein complexes and assemblies, which also had a long history of development but did not progress as fast as the tertiary structure prediction (Lensink et al., 2016, 2018, 2021). However, the situation started to change as more and more deep learning methods were developed to predict inter-protein contacts and quaternary structures(Quadir et al., 2021; Xie & Xu, 2021; Yan & Huang, 2021; Evans et al., 2022; Guo et al., 2022; Roy et al., 2022). Particularly, adapting AlphaFold2 for protein quaternary structure prediction (i.e., AlphaFold-multimer (Evans et al., 2022)) drastically improved the accuracy of protein complex/assembly structure prediction.

In parallel to the development of the methods for sampling quaternary structures, numerous methods have been developed for estimating the accuracy of predicted quaternary structural models (**EMA**) in order to select the best predicted structures and use them effectively in biological research(Dapkūnas et al., 2021). The EMA methods can be classified into four categories: (1) single-model energy/statistical potential-based methods; (2) single-model machine learning methods; (3) multi-model consensus methods; and (4) hybrid methods of combining (1), (2) and/or (3). Single-model energy/statistical potential-based methods (Chen et al., 2003; Lu et al., 2003; Pierce & Weng, 2007, 2008) calculates the physical energy (e.g., electrostatics, hydrophobic interactions, solvation) or statistical potential(Olechnovič & Venclovas, 2017) (e.g., residue-residue contact potentials) of a model to estimate its quality. Machine learning methods including deep learning methods (Wang et al., 2020; Igashov et al., 2021; Wang et al., 2021; Chen et al., 2022; Liu et al., 2023) use expert-curated or automatically extracted features of a model as input to predict its quality. For instance, DProQA (Chen et al., 2022) uses a gated-graph transformer to take a protein quaternary structure represented as a graph (nodes: residues and edges: residue-residue interactions) as input to predict its quality score. The multi-model consensus methods leverage the similarity between structural models to evaluate their quality, which have been extensively used in assessing the quality of protein tertiary structures (Zhang & Skolnick, 2004a; Wallner & Elofsson, 2006; Larsson et al., 2009; Cheng et al., 2009; Wang et al., 2011; McGuffin et al., 2021) and is considered one of the most effective methods for tertiary structure quality assessment(Cheng et al., 2019). The reason that the model consensus methods work well in many situations is that the good models similar to the native structures must be similar to each other, while bad models can have many ways to be incorrect and are therefore often much less similar to each other, leading to a relatively higher average pairwise similarity score for good models. The approach generally works well when there is a significant or relatively high proportion of good models in a model pool that dominate over the bad models. However, the model consensus approach was rarely used in estimating the accuracy of protein *quaternary* structures until the 2022 CASP15 experiment.

Before the CASP15 experiment, we extended our pairwise model similarity method for *tertiary* structure quality assessment (Wang et al., 2011) to estimate the quality of protein *quaternary* structural models. However, the average pairwise similarity between models cannot work well when many models in a model pool have low quality and are also similar to each other (e.g., structure predictors making the same rather than different folding mistakes), leading to bad models having high pairwise similarity scores (i.e., higher estimated quality scores) to be ranked higher. To address the weakness, we developed a hybrid method (MULTICOM_qa) combining the pairwise similarity score between a multimer model and other models (PSS) with a single-model-based interface contact probability score (ICPS) that does not depend on the similarity between models to estimate the accuracy of the model. ICPS is calculated from the inter-chain contact map predicted by inter-chain residue-residue contact/distance predictors (Guo et al., 2022; Roy et al., 2022; Xie & Xu, 2022) such as CDPred (Guo et al., 2022). MULTICOM_qa uses a weighted average of PSS and ICPS to predict the global accuracy (quality) of a quaternary structural model. We first benchmarked MULTICOM_qa on the CASP14 dataset and then blindly tested it in the 2022 CASP15 experiment. It ranked first out of 24 EMA predictors in estimating the global fold accuracy of protein quaternary structural models in CASP15. Our post-CASP15 analysis investigates several important factors influencing MULTICOM_qa’s performance and identifies its strengths and weaknesses for further improvement.

## 2. Methods

Before CASP15, we developed several individual EMA methods to measure the pairwise similarity between multiple quaternary structural models and predict the quality of the contact interfaces of a single structural model, which are described in Section 2.1. We then combined them to design a hybrid method of estimating model accuracy for the CASP15 experiment, which is described in Sections 2.2 and 2.3.

## 2.1 Individual EMA methods

### Average pairwise similarity score (PSS)

A complex structure alignment tool - MMAlign (Mukherjee & Zhang, 2009) - is used to calculate the structural similarity score (i.e., TM-score(Zhang & Skolnick, 2004b)) between a multimer model and each of the other multimer models in a model pool for a multimer target. The average TM-score between a model and all other models is the average pairwise similarity score (**PSS**) for the model (i.e., 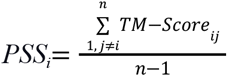, *PSS_i_* is PSS of model *i*, *j* is the index of any model other than *i*, *TM-score_ij_* is the TM-score between model *i* and model *j*, and *n* is the total number of models). PSS is considered an estimation of the true quality of a model. As a comparison for analyzing the effectiveness of the PSS, the average similarity between the monomer structural units of one multimer model and their counterparts of each of the other multimer models (called *monomer_pss*) and the average similarity between the pairs of chains (dimers) of one multimer model and their counterparts of each of the other multimer models (called *dimer_pss*) are also calculated. Monomer_pss and dimer_pss are calculated by TM-score (Zhang & Skolnick, 2004b) and DockQ(Basu & Wallner, 2016), respectively. DockQ measures the similarity between two dimer structures considering multiple factors such as RMSD of two dimer structures, RMSD of one unit (ligand) after another unit (receptor) is superimposed, and the percent of common inter-chain residue-residue contacts. Different from monomer_pss considering only tertiary structure similarity but not inter-chain interactions and dimer_pss considering both tertiary structure similarity and dimeric interactions, PSS assesses the overall fold and interface similarity between a multimer model and all the other multimer models and therefore is expected to be more informative about the quality of the model than mnomer_pss and dimer_pss. It is worth noting that PSS, monomer_pss and dimer_pss are *multi-model* EMA methods because multiple structural models for the same target are required to compute them.

### Average interface contact probability score (ICPS)

The inter-chain residue-residue contacts in a multimer model are first identified. Two residues are considered in contact if their Ca - Ca atom distance is less than 8 Å. The interaction interfaces between two chains that interact in at least 20% of multimers are selected for the ICPS calculation. The inter-chain residue-residue contact probability map for every two chains forming an identified interaction interface are then predicted by an inter-chain residue-residue contact predictor - CDPred(Guo et al., 2022). CDPred is a deep learning-based tool that can predict the inter-chain distance map for homodimers and heterodimers. It leverages the 2D attention mechanism and deep residual network to capture the interaction between different chains to increase the accuracy of inter-chain distance prediction. The inter-chain distance map predicted by CDPred is converted into an inter-chain contact probability map. The predicted probabilities for the inter-chain residue-residue contacts in each interaction interface in a multimer model are extracted from the inter-chain contact probability maps predicted by CDPred. The average probability of the inter-chain residue-residue contacts in the interaction interfaces is called the average interface contact probability score (**ICPS**), measuring the likelihood of the interaction interfaces in a multimer model. For example, **Figure 1** illustrates how ICPS is calculated for an interface. ICPS is a *single-model* quality score because it only needs the information of a single model to be computed, which is different from PSS that depends on the comparison between models.

**Figure 1:**
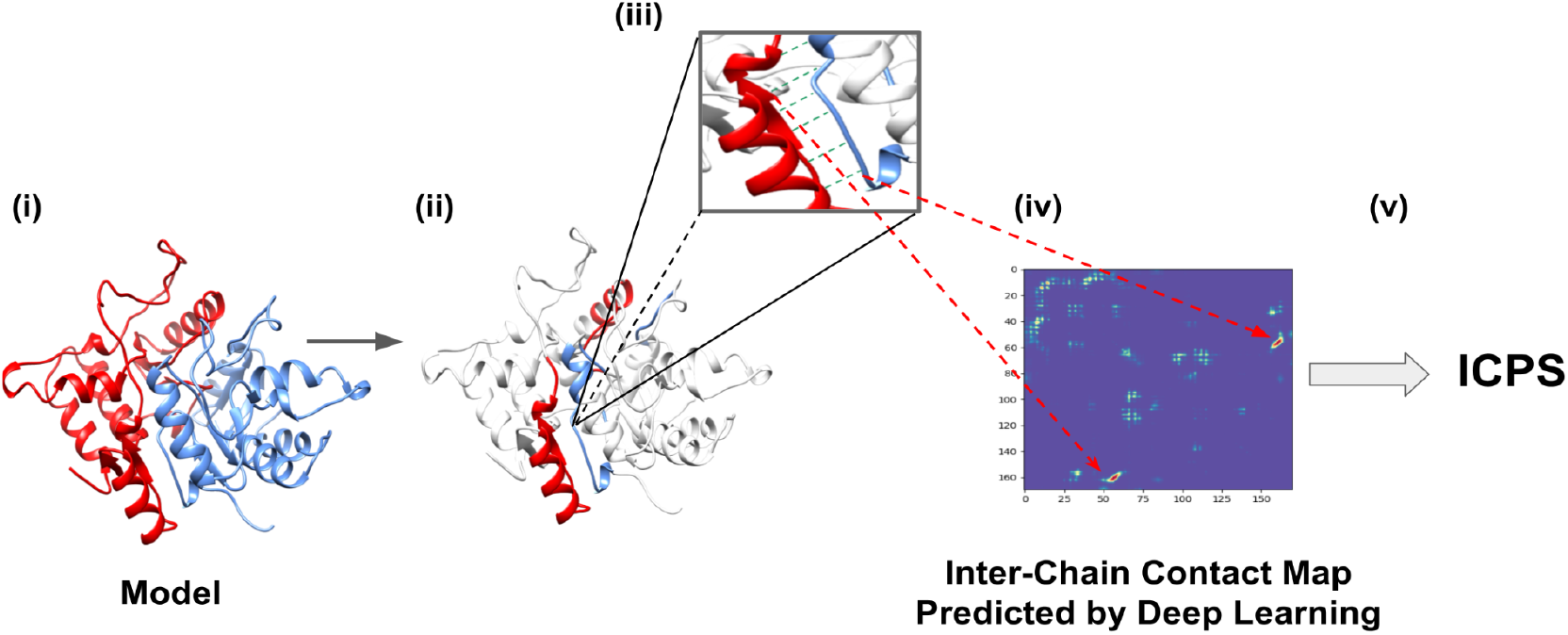
A simplified illustration of the ICPS calculation. **(i)** A dimer model with chain A in red and chain B in blue. **(ii)** All the inter-chain contacts from a dimeric interface of the model are identified, which are coloured in red and blue. **(iii)** The green lines highlight some inter-chain contacts present in the interface. **(iv)** The predicted probability scores for the inter-chain contacts are extracted from the inter-chain contact map predicted by a deep learning predictor (CDPred). **(v)** The probability scores of the contacts in the interface are averaged as ICPS for the model.

### Number of interface contacts (interface_size)

A simple metric - the number of inter-chain contacts in a multimer, which measures the size of the interaction interfaces of a multimer model, was shown to have some discriminative power of ranking multimer models in (Bryant et al., 2022). In this study, this method serves as a baseline to measure the performance of PSS, monomer_pss, dimer_pss and ICPS.

We evaluated the five methods above in terms of average ranking loss on the CASP14 dataset. The CASP14 multimer dataset consists of 21 multimer targets, each of which has about 100 structural models predicted by CASP14 assembly predictors. Each method was used to rank the models for each target. The ranking loss is the true TM-score of the best model for a target minus the true TM-score of the no. 1 model ranked by a method. If a method ranks the truly best model of a target no. 1, the ranking loss for the target is 0. The average per-target ranking loss of each method over all the CASP14 targets was computed (supplementary **Figure S1**).

PSS has the lowest loss of 0.118, which is much lower than the other methods, indicating it is quite effective in ranking CASP14 multimer models. interface_size has the highest loss of 0.319. The loss of dimer_pss is 0.147, lower than 0.199 of monomer_pss and 0.204 of ICPS. dimer_pss performs better than monomer_pss because the former considers the interactions between protein chains but the latter does not. It is also interesting to see that the single-model score - ICPS performs similarly to a multi-model score - monomer_pss and much better than the interface_size, indicating that ICPS contains valuable information for ranking multimer structural models.

### 2.2. Combination of EMA Methods

After establishing PSS as the best individual model ranking method on the CASP14 dataset, we tried to combine different methods (multimer_pss, dimer_pss, monomer_pss, and ICPS) to further improve the model ranking performance. In one experiment, we used the average of the four methods ( (PSS + dimer_pss + monomer_pss + ICPS) / 4) to rank the models of the CASP14 targets. However, the average ranking loss of this combination is 0.156, higher than 0.118 of using PSS alone. The reason for the worse performance is that the dimer_pss and monomer_pss are redundant and inferior to PSS and therefore combining them with PSS does not help.

In another experiment, we tested the combination of the multi-model method - PSS and the single-model method - ICPS. We used each of them to rank the models for each CASP14 target. The weighted average of the two ranks for each model (i.e., 0.4 × rank based on ICPS + 0.6 × rank based on PSS) from the two rankings is used as the final rank for the model. The ranking loss of this average ranking approach is 0.105, lower than 0.118 of using only PSS, indicating that ICPS and PSS are complementary and combining them can improve model ranking. Therefore, we chose to combine PSS and ICPS to develop an EMA predictor for the CASP15 experiment. However, the weights of combining them were not fully optimized prior to CASP15 due to the time constraint.

### 2.3 MULTICOM_qa for Estimating Assembly Model Accuracy in CASP15

Based on the limited test on the CASP14 dataset prior to CASP15, we chose to use the combination of PSS and ICPS to build an EMA method - MULTICOM_qa for CASP15. Because CASP15 required an estimated global quality score for each model instead of its predicted rank, MULTICOM_qa used the weighted average of PSS and ICPS (i.e., 0.4 × ICPS + 0.6 × PSS) as the predicted global fold accuracy score for each model. ICPS was also used as the predicted interface score for each model. CDPred was used to predict inter-chain contact maps for the ICPS calculation. The prediction pipeline of MULTICOM_qa is illustrated in **Figure 2**.

**Figure 2.**
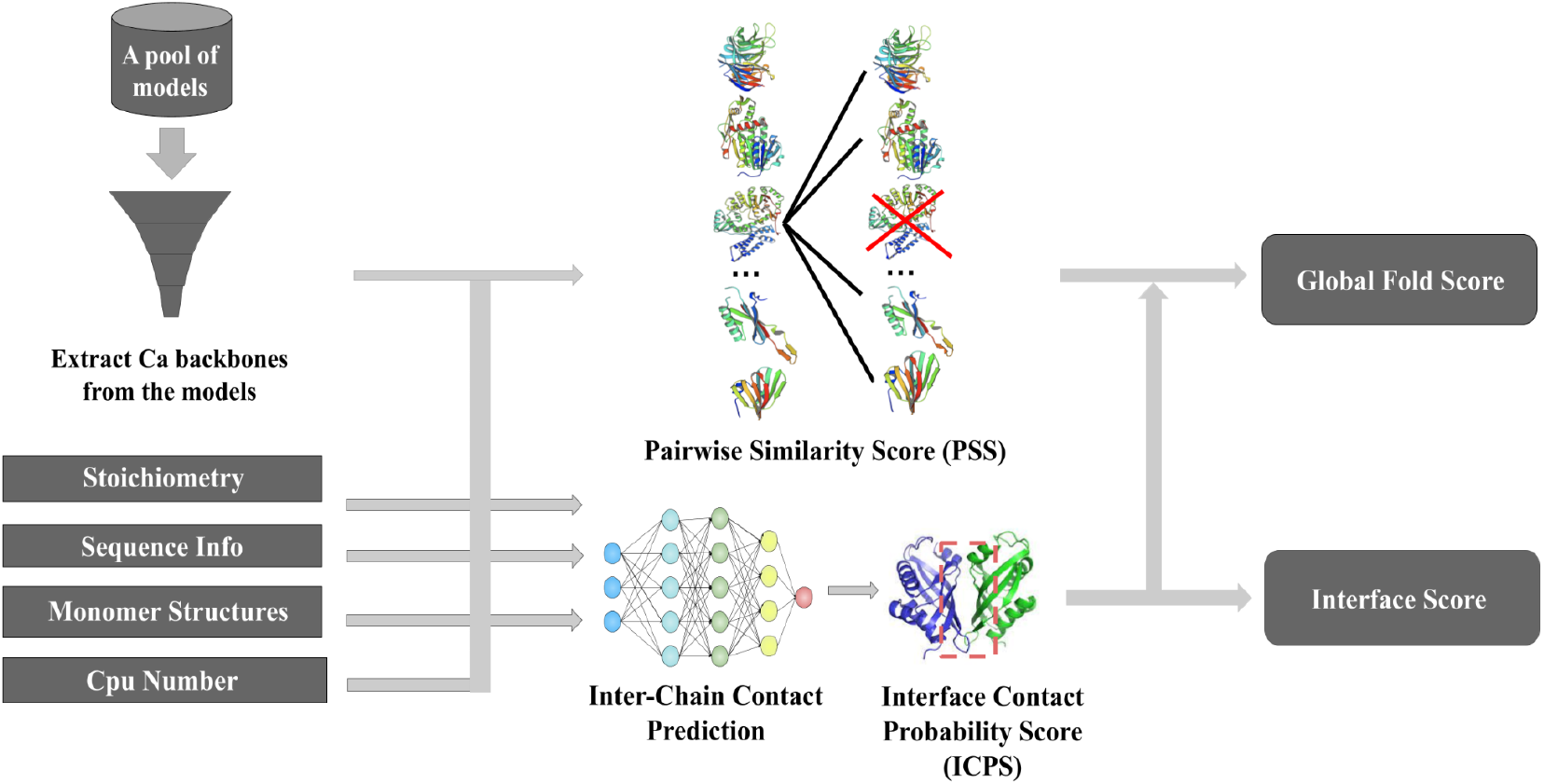
The pipeline of MULTICOM_qa for estimating CASP15 assembly model accuracy. The input is a pool of assembly models for a target and other related information such as the stoichiometry of the target, protein sequences, predicted tertiary structures for each unit in the target, and the number of CPU cores. Multiple CPU cores can be used to speed up the calculation of the pairwise similarity scores between assembly models. PSS and ICPS for each model are computed and then averaged as the predicted global fold accuracy score for the model. The ICPS score is also used as the predicted interface score for each model.

Given a pool of models for an assembly target, MULTICOM_qa extracts their backbone structures by filtering out their side chain atoms. The pairwise similarity score (i.e., TM-score) between any two models is calculated by MMalign. To speed up the calculation, multiple CPU cores can be used to compute the scores in parallel. The average pairwise similarity score between one model and all other models (PSS) is computed for each model. The stoichiometry and sequence information is used to find all the acceptable interactions for a target. A dimeric interaction between two chains is considered as an acceptable interaction and is taken into account in the calculation of the ICPS if it is present in at least 20% of the models in the pool. The tertiary structure for each chain of the target predicted by AlphaFold2 is provided for CDPred to predict the inter-chain residue-residue contact map for any two interacting chains, from which the ICPS for the interfaces of each assembly model is computed. The weighted average of ICPS and PSS for each model is used as the predicted global fold accuracy score of the model, while ICPS is used as the predicted interface score.

MULTICOM_qa was internally benchmarked on the CASP14 dataset with four existing methods: ZRANK(Pierce & Weng, 2007), ZRANK2(Pierce & Weng, 2008), GNN_DOVE(Wang et al., 2021), and DProQA(Chen et al., 2022) (supplementary **Table S1**). MULTICOM_qa clearly performed better than the other four methods. After being briefly tested, MULTICOM_qa blindly participated in the CASP15 experiment from May to August 2022.

## 3. Results

### 3.1 Evaluation metrics and CASP15 assembly targets and models

The CASP15 assessors of the EMA category used a comprehensive evaluation metric (called SCORE) composed of Pearson’s correlation, Spearman’s correlation, ranking loss, and area under ROC curve in terms of both TM-score (Zhang & Skolnick, 2004b) and GDT-TS score (Zemla, 2003) to compare the performance of CASP15 EMA predictors in estimating the global fold accuracy of CASP15 assembly models (Studer et al., 2022). Based on this metric, MULTICOM_qa ranks No. 1 in estimating the global accuracy of the CASP15 assembly models (see the CASP15 official ranking in terms of SCORE: https://predictioncenter.org/casp15/zscores_EMA.cgi for details).

To complement the CASP15 official assessment, in this study, we mainly use two individual, complementary metrics (i.e., *per-target correlation correlation coefficient* (**PTCC**) and *per-target ranking loss* (**PTRL**)) to evaluate the CASP15 global model accuracy prediction results of MULTICOM_qa and analyze its strengths and weaknesses. PTCC is the Pearson’s correlation coefficient between model quality scores predicted by an EMA method (e.g., MULTICOM_qa) and true TM-scores (global quality) of the models of a target. Higher PTCC, the better the estimation of the accuracy of the models of the target. PTRL for a target is the difference between TM-score of the truly best model in a model pool and TM-score of the no. 1 model selected by an EMA method for a target. Smaller PTRL, the better the ranking for the target. PTCC and PTRL measures how well an EMA method performs on one target from the two complementary perspectives. PTCC (or PTRL) can be averaged over all the targets to assess the overall performance of an EMA method on a dataset.

We evaluate MULTICOM_qa’s EMA predictions for the assembly models of all the CASP15 assembly targets except H1171, H1172, and H1185 (i.e., 36 targets in total). H1171 and H1172 with multiple native conformations and H1185 whose native structure was not available to us are excluded. The assembly models for the targets were predicted by CASP15 assembly predictors and were released for CASP15 EMA predictors including MULTICOM_qa to estimate their accuracy from May to August, 2022. Each target has about 275 structural models. The 36 assembly (multimer) targets include 13 homodimers, 10 heterodimers, 6 homomultimers with more than two chains, and 7 heteromultimers with more than two chains. Some targets (e.g., H1111, H1114, T1115, and H1137) are very large. For instance, H1137 has 10 chains with a stoichiometry of

A1B1C1D1E1F1G2H1I1. Almost all the models for these targets were generated by the customized AlphaFold-multimer used by the CASP15 assembly predictors, and most targets have some models of good or reasonable quality. However, quite a few targets are very hard and have few good models, posing a significant challenge for EMA.

### 3.2 The overall performance of estimating the accuracy of CASP15 assembly models and the factors influencing the performance

**Figure 3A** shows the distribution of PTCC between MULTICOM_qa predicted global quality scores and the true TM-scores of the models for the 36 CASP15 assembly targets. The true TM-score of a model is obtained by using MMalign to compare it with the native structure and is normalized with respect to the length of the native structure. Higher PTCC indicates the more linear consistency between the predicted global accuracy and the true global accuracy of the models of a target. 26 targets have a relatively high PTCC >= 0.7, indicating MULTICOM_qa can estimate the relative global quality of the models of most targets reasonably well. The average PTCC on the 36 CASP15 targets is 0.6626. Two targets (H1141 and H1111) have a moderate correlation between 0.35 and 0.65, while another three targets (T1160, H1144 and T1176) have a low positive correlation between 0 and 0.2. MULTICOM_qa completely failed on five targets (T1121, T1187, H1142, H1140 and T1161) as the PTCCs for them are negative.

**Figure 3.**
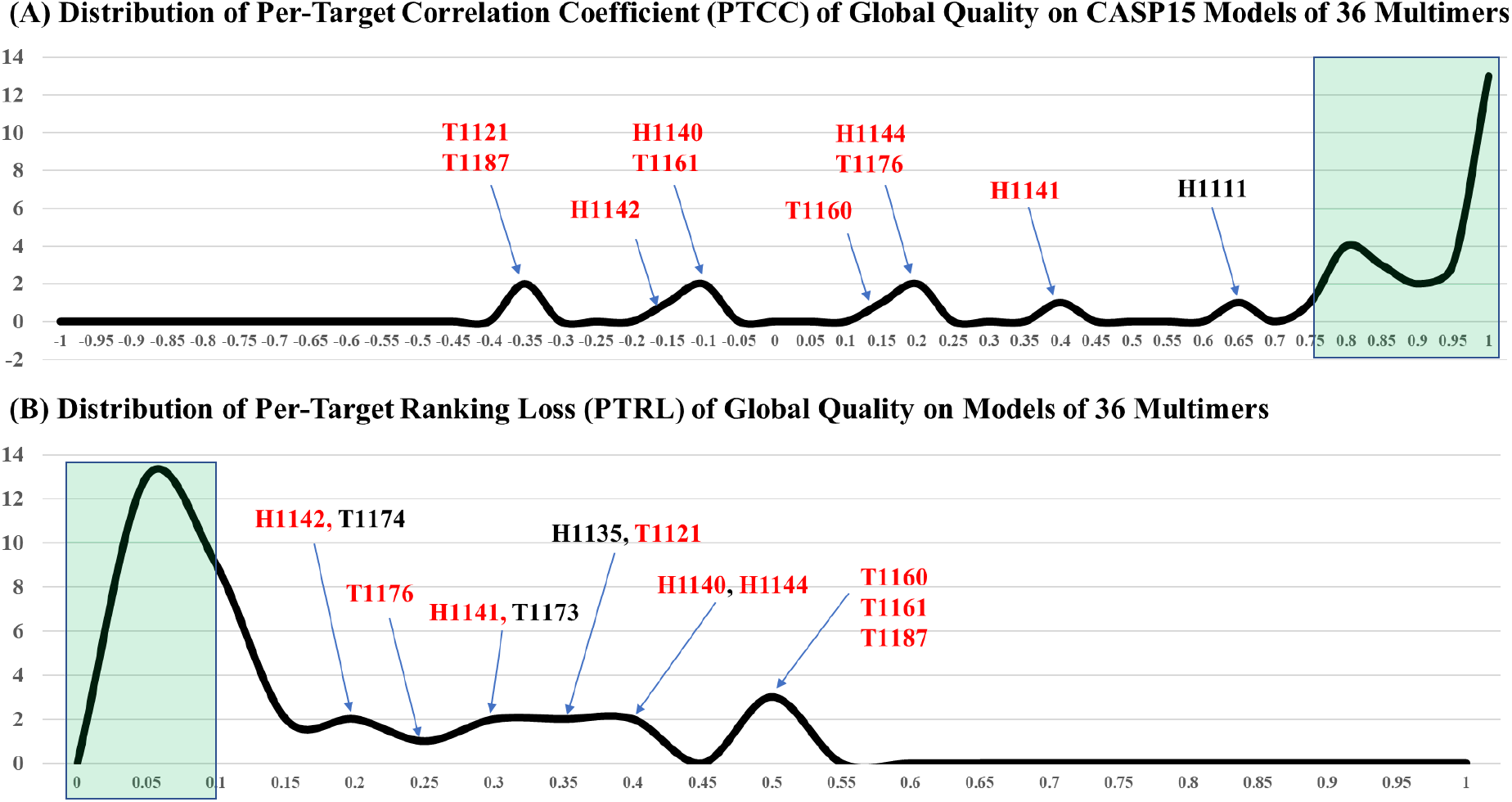
The overall performance of MULTICOM_qa on the models of 36 CASP15 multimer targets. (**A)** Distribution of the per-target correlation coefficient between MULTICOM_qa predicted and true global quality scores (PTCC). The green area contains targets with PTCC > 0.75. Several targets with low/moderate correlation coefficients are identified. The average PTCC on the 36 targets is 0.6626. (**B)** Distribution of the per-target ranking loss (PTRL). The green area contains targets with PTRL <0.1. Several targets with high ranking loss are labelled. The red color highlights nine targets with both high ranking loss and low correlation. The average PTRL on the 36 targets is 0.142. There is a strong negative Pearson’s correlation (−0.8358) between PTCC and PTRL.

**Figure 3B** illustrates the distribution of PTRL for the models of 36 CASP15 multimers. The average PTRL is 0.142. Most targets have a low or acceptable loss less than ~0.15, indicating that MULTICOM_qa can select a reasonable model for most targets as no. 1 model. However, 12 targets (H1142, T1174, T1176, H1141, T1173, H1135, T1121, H1140, H1144, T1160, T1161 and T1187) have a high ranking loss (PTRL > ~0.15), indicating MULTICOM_qa failed to rank good models of these targets at the top. Interestingly, 9 out of the 12 targets (i.e., H1142, T1176, H1141, T1121, H1140, H1144, T1160, T1161 and T1187) also have a low or negative correlation coefficient (PTCC < 0.4), indicating poorly predicted quality scores often lead to a high ranking loss. Indeed, there is a highly negative correlation of −0.84 between PTRL and PTCC (**Figure 4**). Most of the targets that have high PTCC (>0.7) have a relatively low ranking loss (PTRL < 0.15). But there are three pronounced exceptions (T1174, T1173 and H1135) that have high PTCC and high PTRL. Supplementary **Figure S2** plots MULTICOM_qa predicted quality scores against the true quality scores of the models of the three targets, respectively. Interestingly, even though there is an overall strong positive correlation between the predicted quality scores and true quality scores across the entire quality score range, their correlation for good models (true TM-score > 0.6 or 0.7) are negative, indicating that MULTICOM_qa was not able to select better models among the good models and instead it chose some mediocre ones from them as no. 1. There are no targets that have low PTCC (< 0.6) and low PTRL (< 0.1).

**Figure 4.**
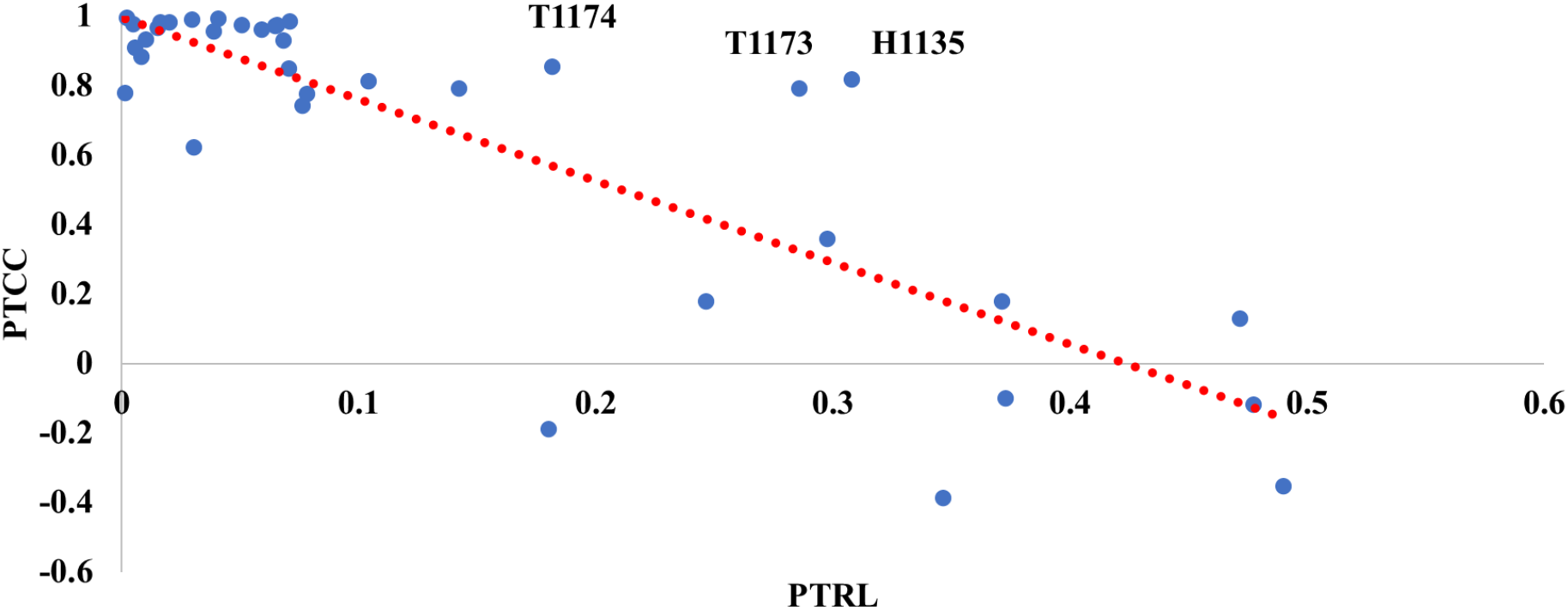
PTCC is plotted against PTRL on the 36 multimer targets. The general trend is that a higher PTCC corresponds to a lower PTRL (correlation between them = −0.8358). There are three pronounced exceptions (T1174, T1173 and H1135) with both high correlation and high ranking loss.

To analyze what factors influence the PTCC of MULTICOM_qa, we plot PTCC against the average true TM-score of the models of each target (i.e., a measure of the absolute difficulty of a target; higher, less difficult), the proportion of good models (the sampling difficulty of a target; higher, less difficult to sample a good model by a method such as AlphaFold-multimer), and the skewness of the distribution of the quality (true TM-scores) of the models of each target (**Figure 5A-C**). The skewness is a measure of the asymmetry of model quality distribution, which is equal to 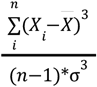, where *n* is the number of the models for a target, *X_i_* is the TM-score of model *i*, 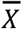 is the mean TM-score and σ is the standard deviation. A large positive skewness indicates that there are more highly concentrated below-average models with a narrow range and there is a wide spread (a long tail) of low-frequency above-average models (supplementary **Figure S3A**). A small negative skewness indicates that there are more highly concentrated above-average models in a narrow range and there is a wide spread (a long tail) of low-frequency below-average models (supplementary **Figure S3B**). A zero skewness means the above-average and below-average models are symmetrically distributed.

**Figure 5.**
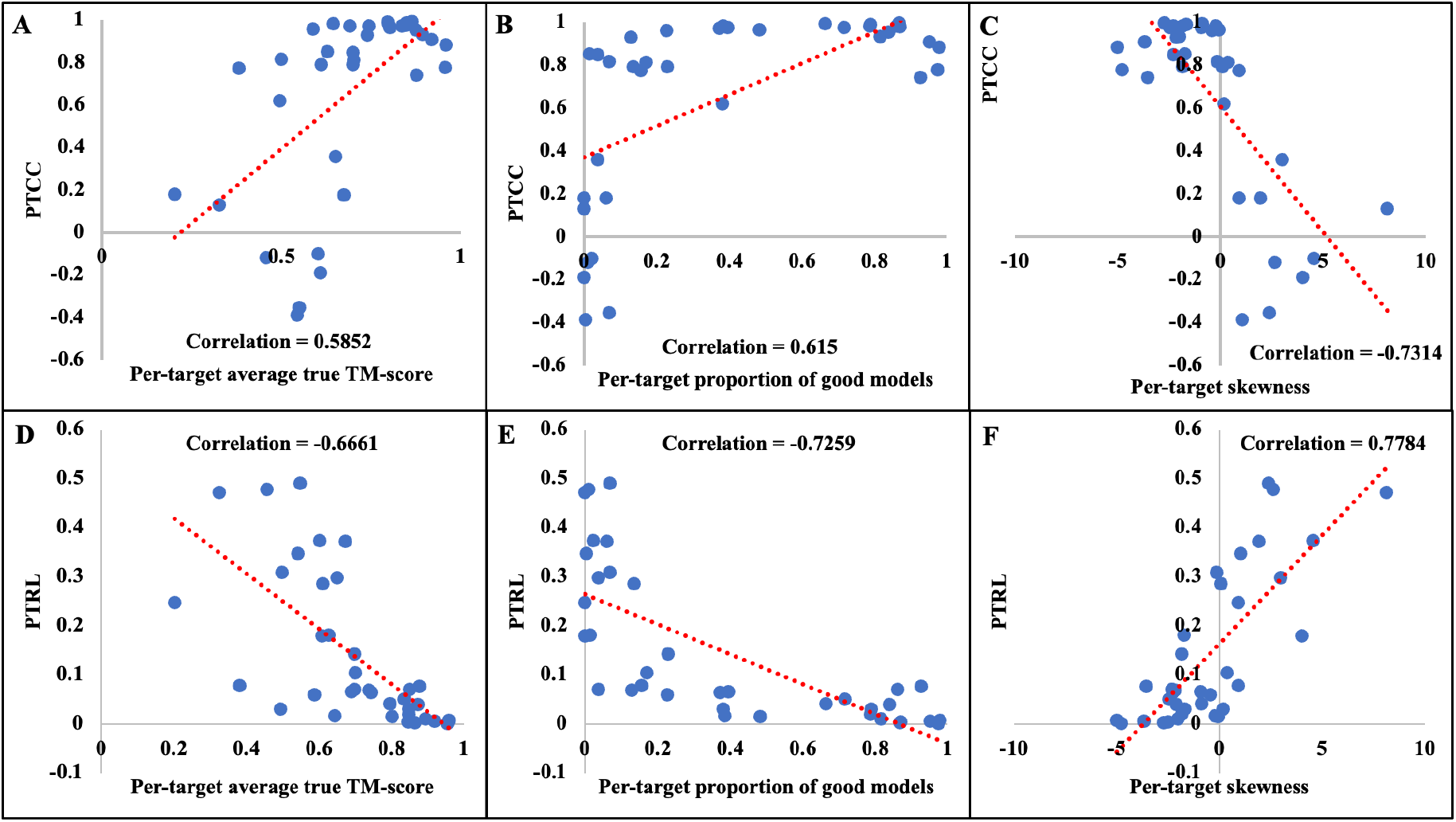
The relationship between PTCC / PTRL of MULTICOM_qa and three factors. **A-C**: Plot of PTCC against **(A)** per-target average true TM-score of the models, **(B)** proportion of good models with TM-score >= 0.8, and **(C)** the skewness of the distribution of the true TM-scores of the models on the CASP15 targets. Each blue dot denotes one target. **D-F**: Plot of PTRL against **(D)** per-target average true TM-score of the models, **(E)** proportion of good models with TM-score >= 0.8, and **(F)** the skewness of the distribution of the true TM-scores of the models on the CASP15 targets.

The Pearson’s correlation between PTCC and each of the three factors (average TM-score, proportion of good models, skewness) is 0.5852, 0.615, and −0.7314, respectively (**Figure 5A-C**). A moderate positive correlation between PTCC and the average TM-score indicates that the absolute difficulty of a target plays a moderate role in estimating the accuracy of the models of the target. A slightly stronger positive correlation between PTCC and the proportion of good models suggests that sampling more good models can make the quality estimation easier, which is expected because more good models lead to higher pairwise similarity scores between good models and likely higher estimated quality scores for them than bad models. A very strong negative correlation between PTCC and skewness indicates that the asymmetry distribution of the quality of the models play an important role in MULTICOM_qa’s estimation of the model accuracy. A high concentration of below-average-quality models in a small range in conjunction with a low frequency of above-average-quality models spreaded in a wide range of quality (high positive skewness) makes the estimation of model accuracy harder. Overall, the skewness influences the model accuracy estimation more than the absolute difficulty of a target (i.e., average TM-score) and the sampling difficulty (i.e., the proportion of good models).

**Figure 5D-F** plots PTRL against three factors (average TM-score, proportion of good models, and skewness). Like PTCC, the similar relationship has been observed that PTRL has a moderate (negative) correlation with the average TM-score measuring absolute target difficulty, stronger (negative) correlation with the proportion of good models measuring the sampling difficulty, and very strong positive correlation with the skewness of the distribution of model quality. For instance, three homomultimers (T1160, T1161 and T1187) with the highest ranking loss of 0.47203, 0.47772 and 0.49026 also have the high skewness of 8.114, 2.6317 and 2.3826.

**Figure 6** compares MULTICOM_qa with other 22 CASP15 EMA predictors (excluding APOLLO since it submitted the same global quality scores for all the models for most targets) in terms of the average PTCC and the average PTRL on 36 CASP15 assembly targets. MULTICOM_qa has the highest average PTCC of 0.6626, which is 17.67% higher than the second highest PTCC of 0.5631 of ModFOLDdock. MULTICOM_qa has the third lowest average PTRL of 0.142, higher than the lowest average PTRL of 0.1067 of Venclovas and second lowest average PTRL of 0.1383 of VoroIF.

**Figure 6.**
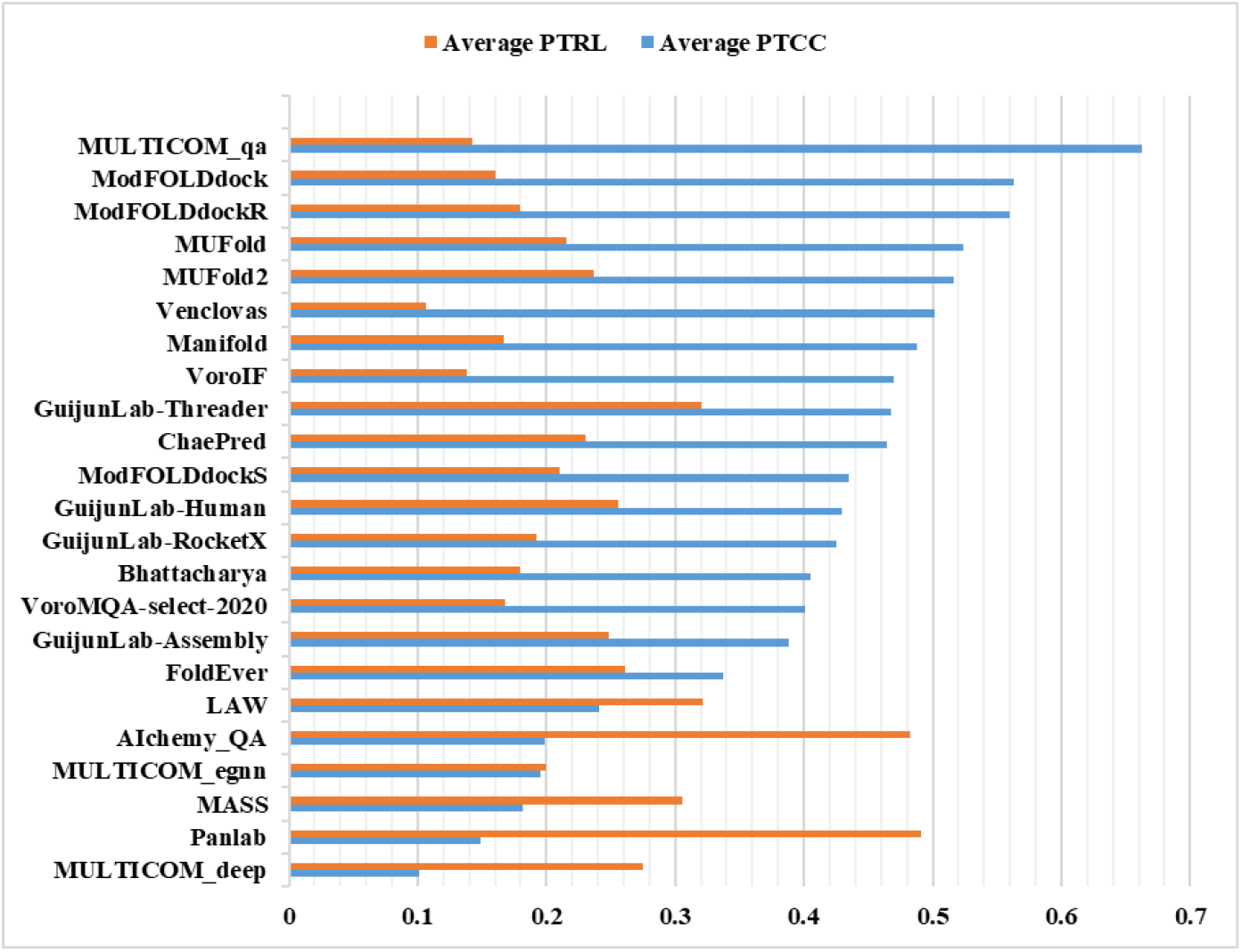
The average PTCC and PTRL of 23 CASP15 EMA predictors. Blue bar denotes the average PTCC. Orange bar denotes the average PTRL. The methods are ordered by average PTCC.

### 3.3 Successful and failed CASP15 cases of MULTCOM_qa

**Figure 7A-D** illustrates the four typical good cases in which MULTICOM_qa performed well and one typical bad case in which it failed. **Figure 7A** illustrates a case (a 27-chain heteromultimer H1111 with stoichiometry of A9B9C9) in which structural models largely fall into two groups: a good group of models with TM-score higher than 0.90 and a bad group of models with TM-score lower than 0.25. Because there is a significant portion of good models and they are highly similar to each other, the average pairwise similarity for good models is relatively high and thus they can be readily picked up by MULTICOM_qa. Indeed, MULTICOM_qa selected one model from the bin with TM-scores in (0.95, 1]. The model has a TM-score of 0.95196, slightly lower than 0.9825 of the best model in the pool (the ranking loss - PTRL = 0.03054). The PTCC between the predicted quality scores and true TM-scores is 0.62, which is only moderate.

**Figure 7.**
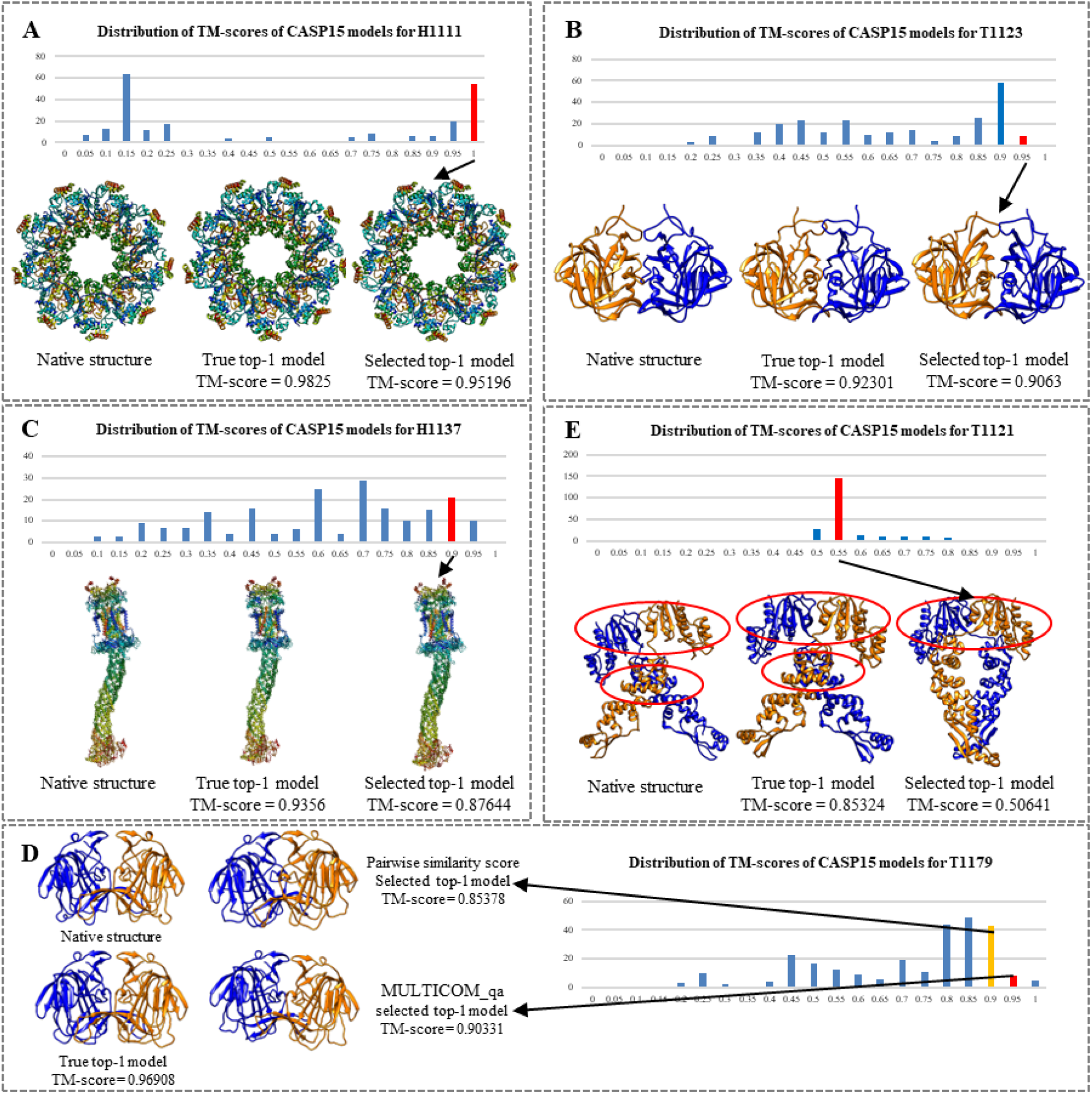
Good and bad examples of applying MULTICOM_qa to estimate the accuracy of the models of CASP15 multimer targets. In each example, the distribution of the true TM-scores of the models for each target is visualized as histogram. The native structure, true top-1 (best) model, selected top-1 model, and the TM-scores of the latter two are presented. The bin in the histogram from which the top-1 model was selected by MULTICOM_qa is highlighted in red. (**A)** H1111: stoichiometry = A9B9C9, PTCC = 0.62, PTRL = 0.03054; (**B**) T1123: stoichiometry = A2, PTCC = 0.98, PTRL = 0.01671; (**C**) H1137: stoichiometry = A1B1C1D1E1F1G2H1I1, PTCC = 0.96, PTRL = 0.05916; (**D**) T1179, stoichiometry = A2, PTCC = 0.97, PTRL = 0.06577; (**E**) T1121: stoichiometry = A2, PTCC = −0.39, PTRL = 0.34683; the correct interfaces are circled; and the top-1 model selected by MULTICOM_qa contains only one of the two correct interfaces, resulting in a high ranking loss of 0.34683.

**Figure 7B** is a homodimer (T1123) that has a large portion of good models with TM-score between 0.85 and 0.9, several high-quality models with TM-score between 0.9 and 0.95, and many models with TM-score widely spanning from ~0.15 to 0.85. MULTICOM_qa was able to select a model from the bin (red) with the highest score range (0.9, 0.95). The top-1 selected model has a TM-score of 0.9063. The PTRL is very low (0.01671). The PTCC is very high (0.98), indicating MULTICOM_qa, mostly its PSS component, is good at estimating the accuracy of the models with this kind of skewed quality distribution.

**Figure 7C** is H1137, a very large heteromultimer target (stoichiometry: A1B1C1D1E1F1G2H1I1). The true TM-score of its models have a wide range of distribution from ~0.05 to 0.95. However, the mode of the distribution lies in the medium quality score range (0.65, 0.7]. MULTICOM_qa was able to select a model from the bin with the second highest score range (0.85, 0.9]. The model has a TM-score of 0.87644, which is not the best but a reasonable choice. The PTRL is 0.05916, which is reasonably low considering the size and difficulty of the target. The PTCC is 0.96, indicating that MULTICOM_qa’s predictions can have high correlation with true quality scores when the true quality scores of the models are distributed in a large range. The situation is somewhat similar to **Figure 7B**.

**Figure 7D** is a case in which combining ICPS with PSS improves the ranking of the models. T1179 is a homodimer whose models have TM-scores largely falling into the range (0.75, 0.9], with some better models on its right side and many worse models on its left side. The pairwise similarity score (PSS) chose a good model with TM-score of 0.85378, but MULTICOM_qa combining PSS and ICPS selected a better model with TM-score of 0.90331. This case indicates that the single-model quality score - ICPS in MULTICOM_qa can sometimes improve the selection of the top-1 model by moving it to the higher quality bin.

**Figure 7E** is a typical case where MULTICOM_qa failed miserably. The quality scores of the models of this homodimer target (T1121) are distributed between ~0.45 and ~0.85. However, the scores of the majority of the models fall into a narrow low score bin (0.50, 0.55], even though there are some better models whose scores are largely evenly distributed from ~0.55 to ~0.85. The distribution of the quality of the models is highly skewed. And because there are too many bad models that are quite similar to each other (average TM-score for the bad models in the (0.5 - 0.55] bin are 0.86), they dominate the calculation of PSS, leading to bad models having higher estimated quality scores. Consequently, MULTICOM_qa chose a bad model from the bin (0.5, 0.55]. The ranking loss is very high (PTRL = 0.34683). Moreover, the PTCC is negative (−0.39), indicating that PSS of MUTLCOM_qa completely failed in this situation and ICPS could not rescue it. This is a typical case in which MULTICOM_qa fails when there is only a small portion of good models and the bad models tend to be similar to each other. Indeed, MULTICOM_qa selected a model with one interaction interface that matches with one of the two interaction interfaces of the native structure, while the best model in the pool has the two correct interfaces.

Supplementary **Figure S4** shows 8 additional good cases where MULTCOM_qa performs well, each of which has a significant portion of good models making ranking easy. Supplementary **Figure S5** shows 6 additional bad cases where MULTICOM_qa failed, each of which has a small portion of good models.

To further analyze how MULTICOM_qa may succeed or fail, we draw a *model similarity graph* for a good case H1111 and a bad case T1121 (**Figure 8**). In each graph, a node denotes a model and an edge connects two models if their structural similarity is higher than a threshold. Here, the threshold is chosen so that 30% of model pairs are connected by edges. The model similarity graph of H1111 has 7 subgraphs (clusters), among which the largest subgraph has higher true quality scores and high within-subgraph similarity. Both the best model and the top-1 model selected by MULTICOM_qa are from the largest subgraph that contains a lot of good and similar models. In contrast, the model similarity graph of T1121 has five subgraphs (clusters). The best model resides in the third large subgraph, while MULTICOM_qa selected one model from the largest subgraph containing a lot of bad models that are similar to each other. This example clearly demonstrates that the high similarity between bad / mediocre models in a large cluster makes model ranking difficult for MULTICOM_qa, mostly its PSS component.

**Figure 8.**
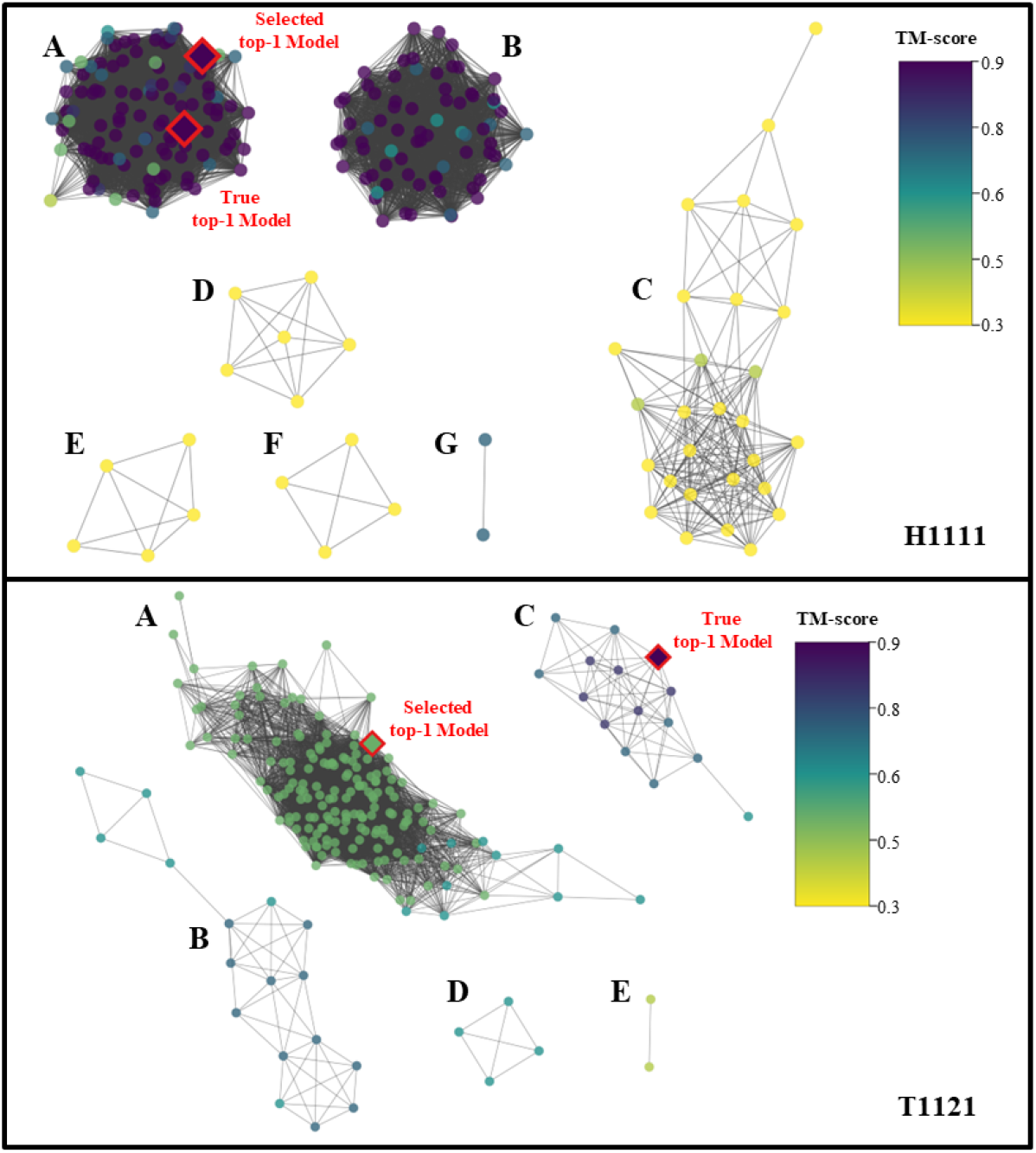
The model similarity graphs for H1111 (top) and T1121 (bottom). Each node denotes a model. An edge is used to connect two models (nodes) if their structural similarity score - TM-score is greater than a threshold. The threshold is determined such that only the top 30% of model pairs are connected by edges. The weight of each edge is TM-score between two nodes (models) calculated by MMalign, which is normalized by the total sequence length of the larger model if the two models have different sizes. The color of the nodes correspond to the true TM-scores of the models. For H1111, both the best model (true top-1 model) and selected top-1 model come from the largest subgraph with the highest quality. For T1121, the selected top-1 model is from the largest subgraph with mediocre quality, but the best model is in the third largest subgraph.

### 3.4 Quantification of the difficulty of structural models for model accuracy estimation

The results in Section 3.2 show that the absolute difficulty of a target (average true TM-score of the models of the target), the sampling difficulty (the proportion of good models), and the skewness of the distribution of quality of the models affect the EMA performance (PTCC and PTRL) of MULTICOM_qa moderately or strongly. We also calculate the correlation between PTCC and the mean pairwise similarity between all the models in a model pool of a target (i.e., equal to the average of PSS of the models in the pool) for the CASP15 targets (**Table 1**). The correlation between MULTICOM_qa’s PTCC and the mean pairwise similarity score of all the models of a target is only 0.1220, which is much weaker than the correlation between PTCC and the other three factors (0.5852 for average TM-score; 0.6150 for the proportion of good models, and −0.7314 for the skewness), indicating that the mean pairwise similarity of all the models in a model pool is not a good indicator of the difficulty of estimating the accuracy of the models. The reason is that, even though high similarity between good models makes the model accuracy estimation easier, high similarity between bad models makes it harder. To treat the negative and positive impact of the similarity between good/bad models differently, we design a new metric called *Difficulty Index* to quantify the difficulty of estimating the accuracy of the models for a target.

**Table 1.** Pearson’s correlation between PTCC (or PTRL) of MULTICOM_qa and each of the five factors (average TM-score, proportion of good models, skewness, mean pairwise similarity score between all the models, and difficulty index). Bold numbers denote the strongest correlation.

| Pearson's Correlation | Avg true TM-score of models | Proportion of good models (TM-score > 0.8) | Skewness | Mean pairwise similarity score between all models | Difficulty Index |
| --- | --- | --- | --- | --- | --- |
| PTCC of MULTICOM_qa | 0.5852 | 0.6150 | -0.7314 | 0.1220 | **0.7723** |
| PTRL of MULTICOM_qa | -0.6661 | -0.7259 | 0.7784 | -0.1478 | **-0.8360** |

The difficulty index for the models of a target is defined as 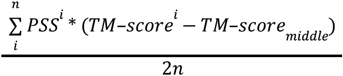. PSS^i^ is the average pairwise similarity score of model i, n is the number of models. 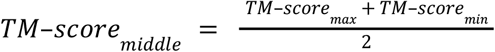. The difficulty index ranges from −1 to 1. Lower the difficulty index, more difficult the models are for accuracy estimation. In this definition, the PSS of the below-average-quality (bad) models whose TM-score is less than median (TM-score_middle_) contributes negatively to the difficulty index, while the above-average-quality (good) models contribute positively. The correlation between the difficulty index and PTCC (or PTRL) of MULTICOM_qa is 0.7723 (or −0.8360), which is even stronger than the correlation between the skewness and PTCC (or PTRL) and several times more stronger than the mean pairwise similarity score between all the models (**Table 1**). The results show that the difficulty index considering the different impacts of the similarity between bad/good models is a very effective measure of the difficulty of estimating the models in a model pool, which is more effective than only considering the skewness of the distribution of the model quality. The correlation for the difficulty index is much higher than that of the mean pairwise model similarity confirms the importance of treating the pairwise similarity between good or bad models differently for EMA. The difficulty index can be used to select hard targets for testing EMA methods in order to improve their performance on them.

Moreover, there is a strong correlation (0.8078) between the difficulty index and the average true TM-score of the models (absolute difficulty) of a target (supplementary **Figure S6**). This indicates that the difficulty index of the models for a target can also quantify the difficulty of predicting structures for the target.

### 3.5 Analysis of the properties and contributions of PSS and ICPS

We compare how the EMA performance (PTCC and PTRL) of the two components of MULTICOM_qa (i.e., PSS and ICPS) is influenced by the four factors (**Table 2**): (1) the average true TM-score of the models of a target, (2) the proportion of good models, (3) the skewness of the distribution of the quality of models, and (4) the difficulty index of the models. The PTCC of PSS has the strongest correlation with the four factors (**Table 2A**), the PTCC of MULTICOM_qa has the second strongest correlation with them (see **Table 1**), and the PTCC of ICPS has a much weaker correlation with them than PSS and MULTICOM_qa (**Table 2A**). This indicates that the performance of the single-model method - ICPS is much less influenced by the four factors than the multi-model method - PSS. The stronger correlation between MULTICOM_qa with the four factors mostly comes from PSS as they have similar correlation coefficients and trends. The largely similar results are observed in terms of PTRL (see the PTRL’s correlation for PSS and ICPS in **Table 2B** and for MULTICOM_qa in **Table 1**). Finally, PTCC/PTRL of PSS has the strongest correlation with the difficulty index, but PTCC/PTRL of ICPS has the strongest correlation with the skewness.

**Table 2.**
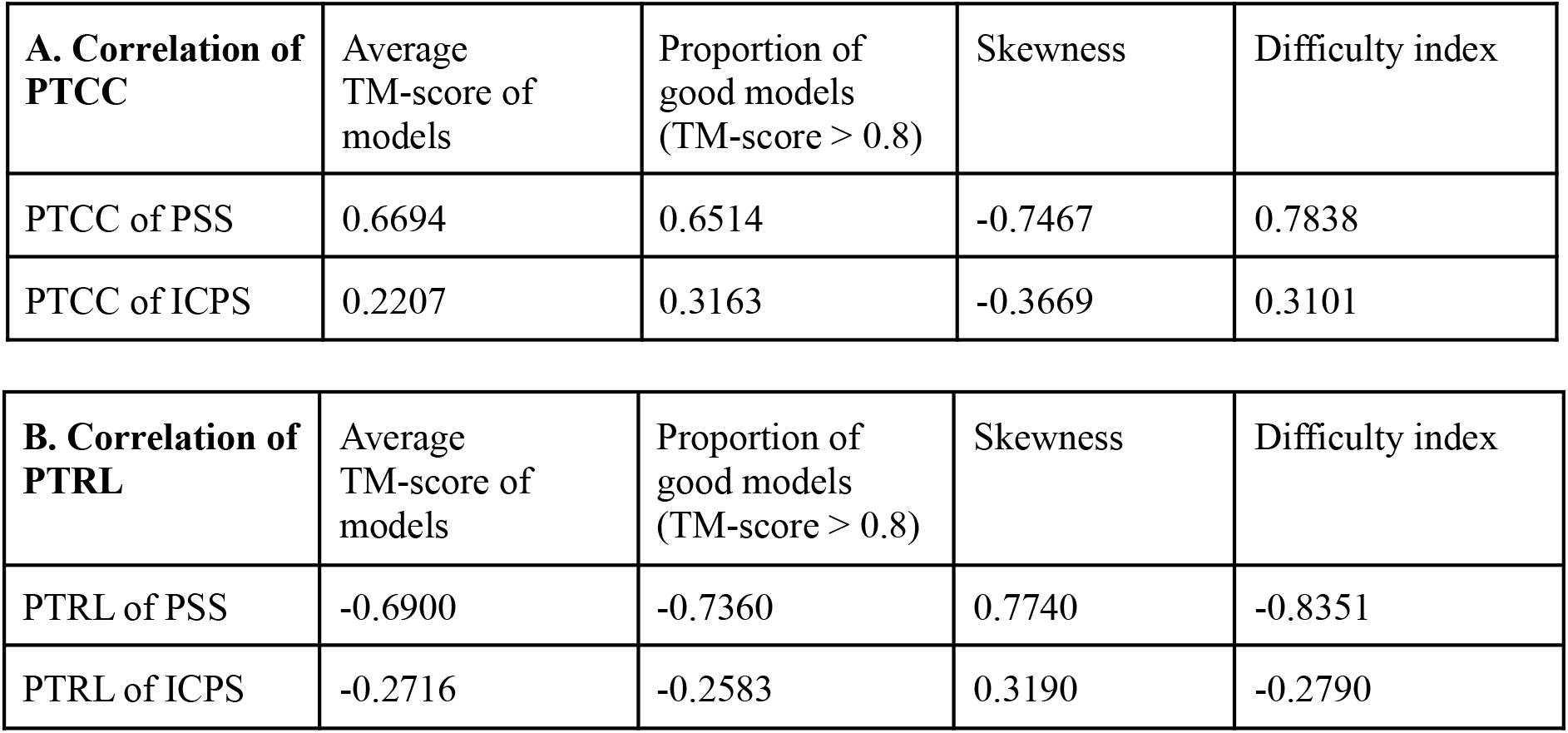
The correlation between the performance (PTCC or PTRL) of two components (PSS and ICPS) of MULTICOM_qa and the four factors (average true TM-scores of the models of a target, the proportion of good models, the skewness of the quality of the models, and the difficulty index of the models). **(A)** The correlation between PTCC and the four factors; and **(B)** the correlation between PTRL and the four factors.

We also investigate how the two components (PSS and ICPS) of MULTICOM_qa contribute to its EMA performance on the 36 CASP15 multimer targets. Because MULTICOM_qa was not able to generate PSS for a very large multimer H1114 ((stoichiometry: A4B8C8)) within the two-day limit, it used ICPS scores alone as the final predicted quality scores for the models of H1114 during the CASP15 experiment. Here, to fairly compare the performance of PSS and ICPS of MULTICOM_qa, we use MMalign (version 2021/8/16) to regenerate PSS for the models of H1114 and calculate the weighted average of the PSS and ICPS as the final predicted scores.

**Table 3** reports the average PTCC and the average PTRL of the three methods on the 36 CASP15 multimer targets. MULTICOM_qa has the lowest average PTRL, while PSS has the highest average PTCC, indicating that combining PSS and ICPS in MULTICOM_qa reduces the average ranking loss, but slightly decreases the average per-target correlation coefficient. Both MULTICOM_qa and PSS perform much better than ICPS, suggesting that the multi-model based method such as PSS still works much better than the single-model based interface contact score - ICPS on average. However, PSS and ICPS are complementary. H1114 (stoichiometry: A4B8C8) is a notable example showing that combining ICPS and PSS reduces the ranking loss. **Figure 9A** illustrates how ICPS and MULTICOM_qa selected a model for H1114 that is much better than PSS. The reason is that CDPred predicted high probabilities for some true contacts in an interface of some good models, leading to a higher ICPS for them to be selected (**Figure 9B**)

**Table 3.** The PTCC and PTRL of PSS, ICPS and MULTICOM_qa (correction) on 36 CASP15 multimer targets. MULITICOM_qa (correction) differs from the CASP15 MULTICOM_qa only on one target - H1114. Bold number denotes the best PTCC or PTRL.

| Method | Average PTCC↑ | Average PTRL↓ |
| --- | --- | --- |
| PSS | **0.6737** | 0.1454 |
| ICPS | 0.3139 | 0.2938 |
| MULTICOM_qa (correction) | 0.6645 | **0.1413** |

**Figure 9.**
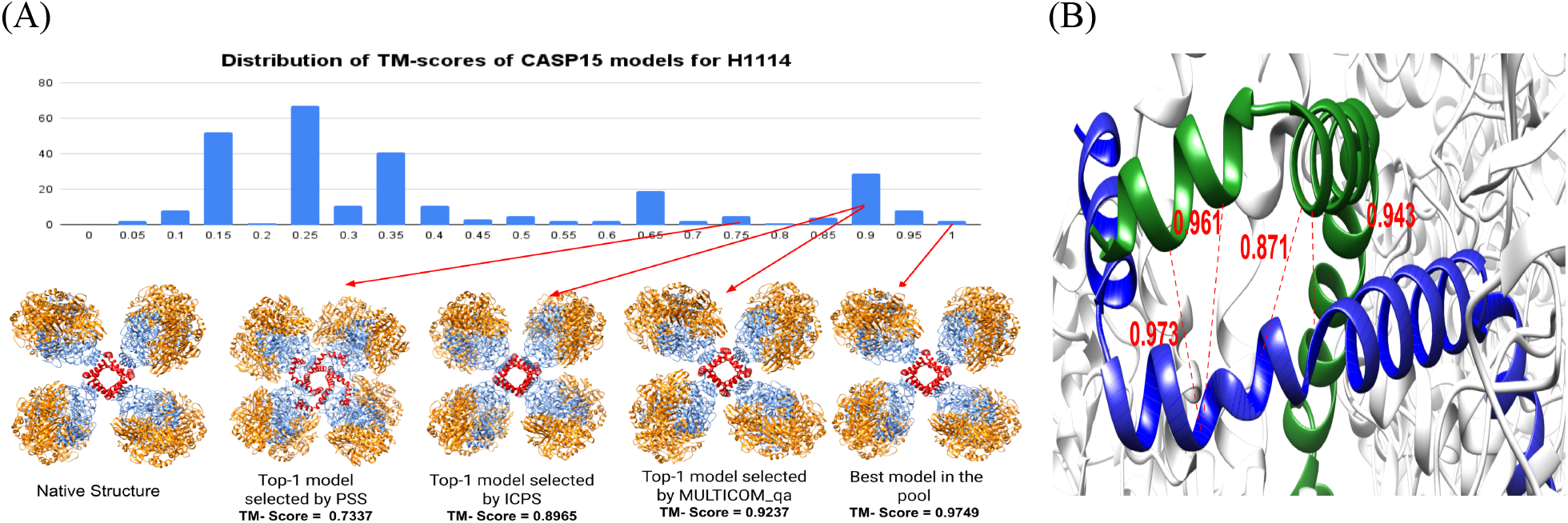
**(A)** The distribution of TM-scores of the models of H1114 as well as its native structure, top-1 model selected by PSS, top-1 model selected by ICPS, top-1 model selected by MULTICOM_qa, and the best model in the model pool. Both ICPS and MULTICOM_qa selected a model that is much better than the model selected by PSS. The four A chains in the good models form a cube in the center (red), which is a key feature of the structure of H1114. **(B)** A homo-dimeric interface between two A chains of the top-1 model selected by ICPS for target H1114. The model is H1114TS119_2. The red lines represent the true inter-chain contacts in the interface. The numbers are the probabilities for the true contacts predicted by the CDPred. The true contacts have high predicted probabilities, leading to a high ICPS of 0.44 for the interface.

### 3.6 Optimization of the weights of ICPS and PSS in MULTICOM_qa

When we developed MULTICOM_qa for the CASP15 experiment, the weights for its two components (0.4 for ICPS and 0.6 for PSS) were not optimized due to the time constraint. After the CASP15 experiment, we tested different weights for them on the 36 multimer targets to see if better results can be obtained. As in Section 3.5, the incorrect PSS of the models of H1114 was corrected for this analysis. **Figure S7** shows how the average per-target ranking loss (PTRL) of MULTICOM_qa changes with respect to the weight of ICPS. The lowest average PTRL of 0.1381 is achieved at the ICPS weight of 0.48, which is better than the loss of 0.1413 for weight 0.4 used in the CASP15 experiment and the loss of 0.1454 of using PSS alone (i.e., ICPS weight = 0). The results show that the performance of combining PSS and ICPS may be further improved if their weights are optimized.

## 4. Discussion

Calculating the similarity between two multimers depends on multimer structure alignment tools such as MMalign. The existing multimer alignment tools require a good (or optimal) mapping between the chains of one multimer and those of another multimer in order to accurately calculate their similarity. However, finding an optimal mapping for two large homomultimers or large heteromultimers containing multiple copies of the same chain is extremely hard and time consuming. Therefore, even though the empirical approach adopted by MMalign can calculate multimer similarity well in most cases, sometimes it severely underestimates the similarity between the models of some large multimers consisting of many chains or yields different similarity scores when the order of the two multimer models is changed, which negatively affects the performance of MULTICOM_qa. Moreover, the speed of calculating similarity between large multimer models consisting of dozens of chains is slow, which prevents the pairwise similarity calculation from being applied to a large number of models. However, it appears that a new, efficient algorithm of finding the optimal chain mapping between large multimers was designed by CASP15 EMA assessors during the CASP15 meeting(Studer et al., 2022), which is expected to alleviate the problem. Given an optimal chain mapping as input, some complex alignment tools such as DockQ that requires a chain mapping as input, can be more readily applied to aligning mutimers consisting of more than two chains to calculate their similarity.

As demonstrated in our CASP15 experiment, the multi-model pairwise similarity approach works well when the quality scores of the models for a target are either shifted toward the high score range or evenly distributed across the score range. On average the pairwise similarity approach (i.e., PSS) still works much better than single-model approaches (e.g., ICPS) of estimating the quality of each model. This is not surprising because the multi-model pairwise similarity is actually measuring the frequency of models sampled by the underlying prediction methods. If the underlying prediction methods do a reasonable job, it is expected to produce good models with high probability. Because good models must be similar to the same native/true structure, the good models must be similar to each other, leading to a higher average pairwise similarity score for them. Another implicit assumption for the pairwise similarity method to work well is that bad models have different ways to be wrong and therefore are likely different to each other. It is true if the underlying prediction methods do not make the same mistake frequently. However, this assumption can be violated, reflected in that the pairwise similarity approach often fails when there is a large portion of bad models that are similar to each other, which dominates the average pairwise similarity calculation so that bad models are assigned higher scores than good models. This happens for some hard targets when the underlying structure prediction methods generate similar bad models for them. This is a key issue to be addressed in order to further advance the pairwise similarity approach. The difficulty of such hard targets can be measured well by the difficulty index introduced in this work.

One way to correct the weakness of the pairwise similarity approach is to combine it with the complementary single-model EMA methods such as the deep learning predicted interface contact probability score (ICPS) that do not depend on the comparison between models. This direction may be important for advancing the state of the art of estimating model accuracy because it is unlikely single-model EMA methods alone can catch up with the performance of the pairwise model similarity approach in a short period of time. However, although this direction is promising, the combination of the two is not trivial. A simple weighted average of the scores of the two can correct some mistakes but may not significantly improve the model accuracy estimation. Therefore, more sophisticated approaches of integrating the strengths of the two complementary approaches need to be developed. For instance, a deep learning method can be trained to combine the pairwise similarity scores with multiple single-model EMA methods to estimate model accuracy. Moreover, for AlphaFold2 predicted multimer models, the AlphaFold2 predicted TM-score or plDDT score can potentially be combined with the pairwise similarity approach if they are available. We have noticed that the model ranking of AlphaFold2 predicted quality score and the pairwise model similarity score are complementary on our in-house models generated for CASP15 targets (data not shown). However, the search for the best approach of combining them is challenging and still ongoing. Moreover, to further advance the combination, more sophisticated and accurate deep learning single-model EMA methods for quaternary structure models need to be developed as what happened in estimating the accuracy of tertiary structure models(Chen et al., 2023).

## 5. Conclusion

In this work, we developed a hybrid method of combining the pairwise model similarity score and the interface residue-residue contact prediction to estimate the accuracy of protein assembly models and blindly tested it in the CASP15 experiment. The method is effective in ranking protein assembly models and predicting their global quality scores as demonstrated by its outstanding performance in the CASP15 experiment. Our experiment demonstrates that the average pairwise similarity score (PSS) can estimate the accuracy of models well in most cases, but it fails when a hard target has a large portion of bad models that are similar to each other. The single-model based interface contact probability score (ICPS) derived from the deep learning-based inter-chain contact prediction can provide some valuable information complementary with PSS to evaluate the quality of the inter-chain interaction interface in multimer models. A simple weighted combination of PSS and ICPS can correct some model ranking errors of PSS in some cases but does not systematically improve its ranking, suggesting that a more sophisticated (e.g., deep learning-based) integration of the pairwise similarity scores and single-model quality scores is needed to significantly improve the estimation of quaternary model accuracy. More accurate single-model quality assessment methods are also needed to improve the combination. Finally, it is useful to consider several important factors such as the target difficulty, model sampling difficulty, skewness of model quality, and model difficulty index for designing and evaluating EMA methods.

## Acknowledgement

Research reported in this publication was supported in part by two NIH grants [R01GM093123 and R01GM146340], Department of Energy grants [DE-SC0020400 and DE-SC0021303], and two NSF grants [DBI1759934 and IIS1763246]. We thank CASP15 organizers and assessors for providing valuable data and advice for this work.

## Authors’ Contributions

JC conceived and designed MULTICOM_qa. RR implemented the method and tested it on the CASP14 dataset. ZG adapted CDPred for the ICPS calculation. RR and NG ran MULTICOM_qa in the CASP15 experiment and collected the CASP15 data. JC, JL and RR designed data analysis methods. JL, JC and RR analyzed the CASP15 results. JC, RR, and JL wrote the manuscript.

## Supplementary Materials

**Figure S1:**
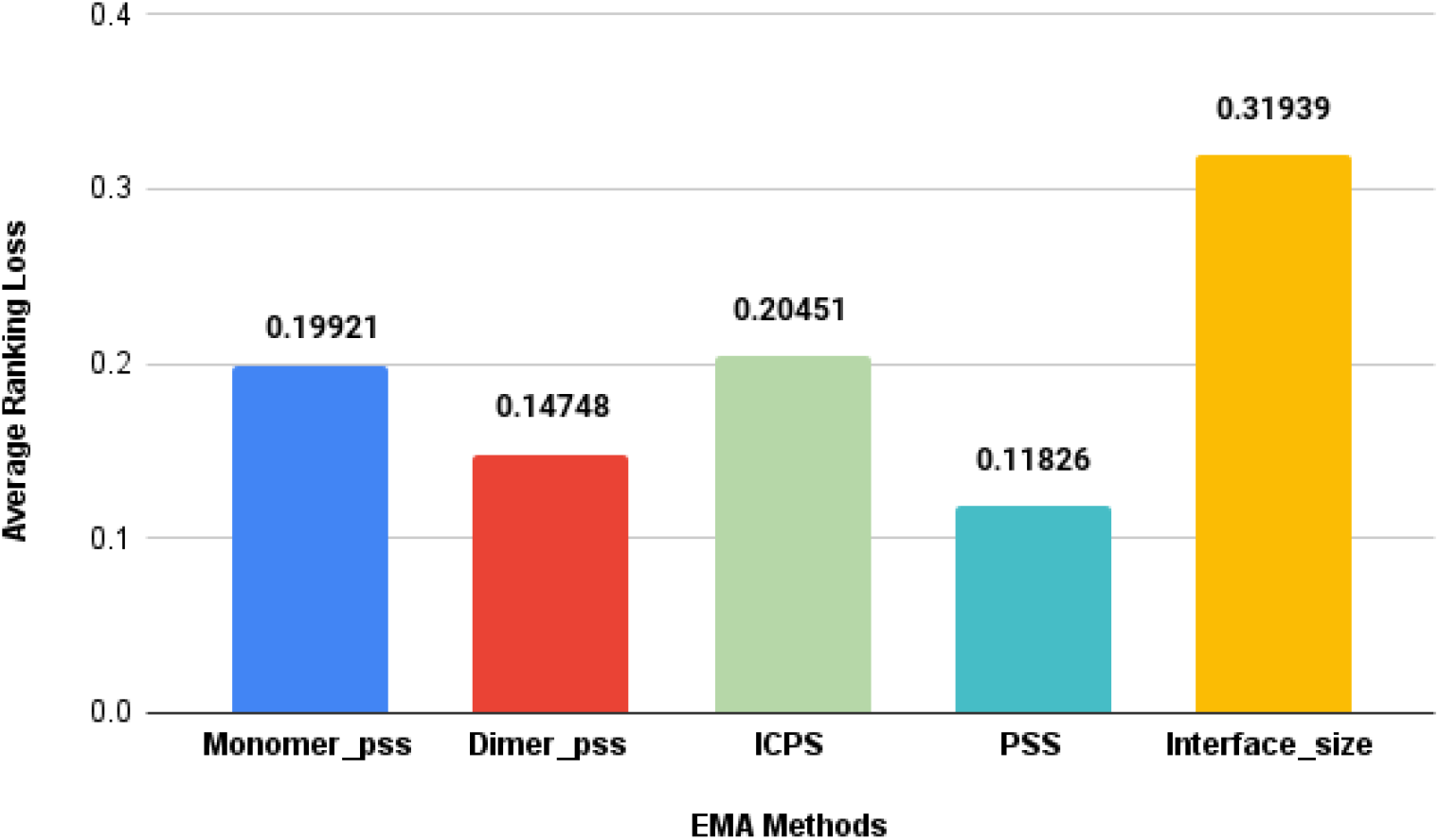
The average ranking loss of the different EMA methods on the CASP14 dataset. PSS has the lowest average ranking loss.

**Table S1.** Average true TM-Score and the average ranking loss of top 1 or the best of top five models selected by each method on the CASP14 dataset. Bold numbers denote the best scores. MULTICOM_qa performs best in terms of the two metrics.

| Method | Average TM-Score↑ |  | Average Ranking Loss↓ |  |
| --- | --- | --- | --- | --- |
|  | top 1 | best of top 5 | top 1 | best of top 5 |
| MULTICOM_qa | **0.539** | **0.598** | **0.146** | **0.0869** |
| ZRANK | 0.385 | 0.5358 | 0.2998 | 0.1486 |
| ZRANK2 | 0.406 | 0.5116 | 0.2781 | 0.1728 |
| GNN_DOVE | 0.338 | 0.463 | 0.3460 | 0.221 |
| DProQA | 0.459 | 0.559 | 0.226 | 0.126 |

**Figure S2.**
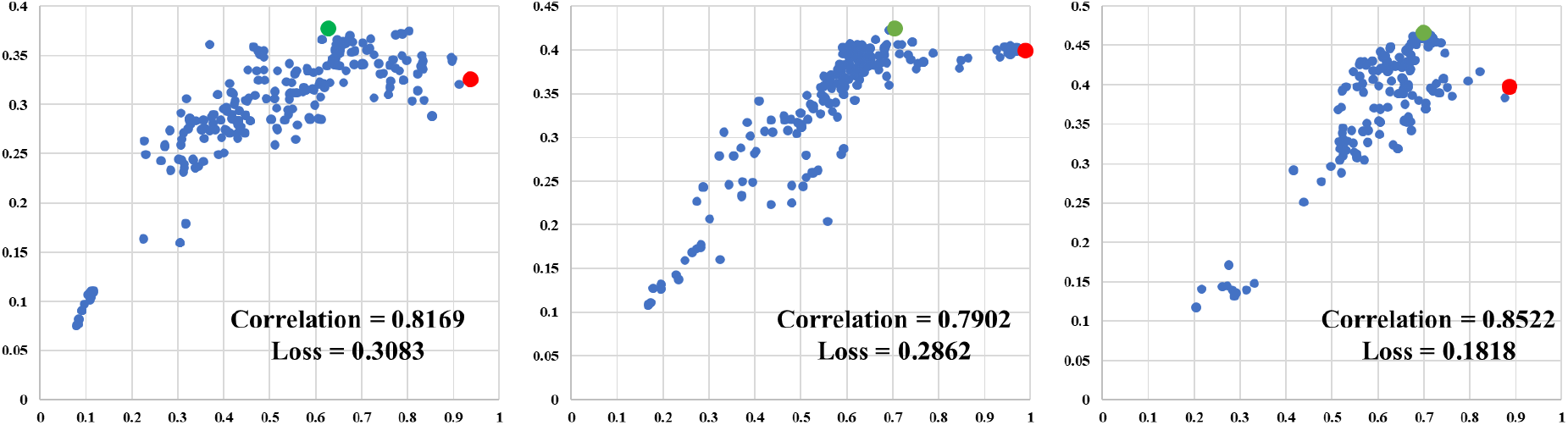
The quality scores predicted by MULTICOM_qa on y-axis are plotted against the true TM-scores on x-axis for the models of H1135 (left, skewness = −0.1433), T1173 (middle, skewness = 0.100) and T1174 (right, skewness = −1.7264). The green dots represent the model with the highest predicted quality scores and the red dots represent the model with the highest true TM-score.

**Figure S3.**
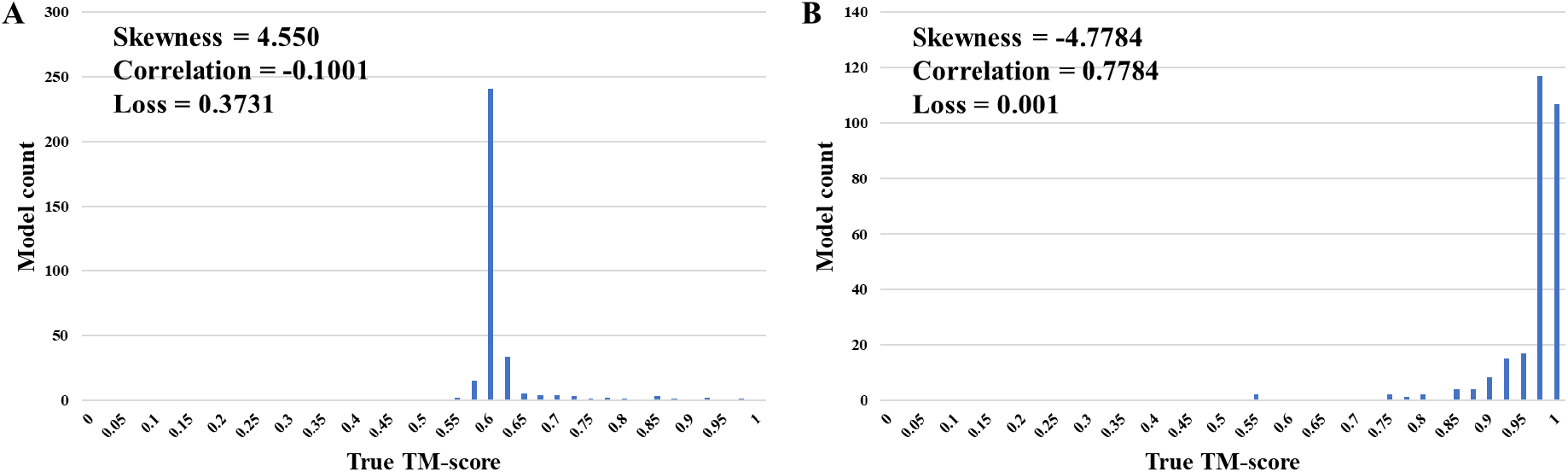
The skewness of the distribution of the quality (TM-score) of the models of a target. **(A)** An example of high positive skewness (H1140); most models have relatively low quality, whereas a small number of models have much higher quality. **(B)** An example of very low negative skewness T1127; most models have relatively high quality, whereas a small number of models have much lower quality.

**Figure S4.**
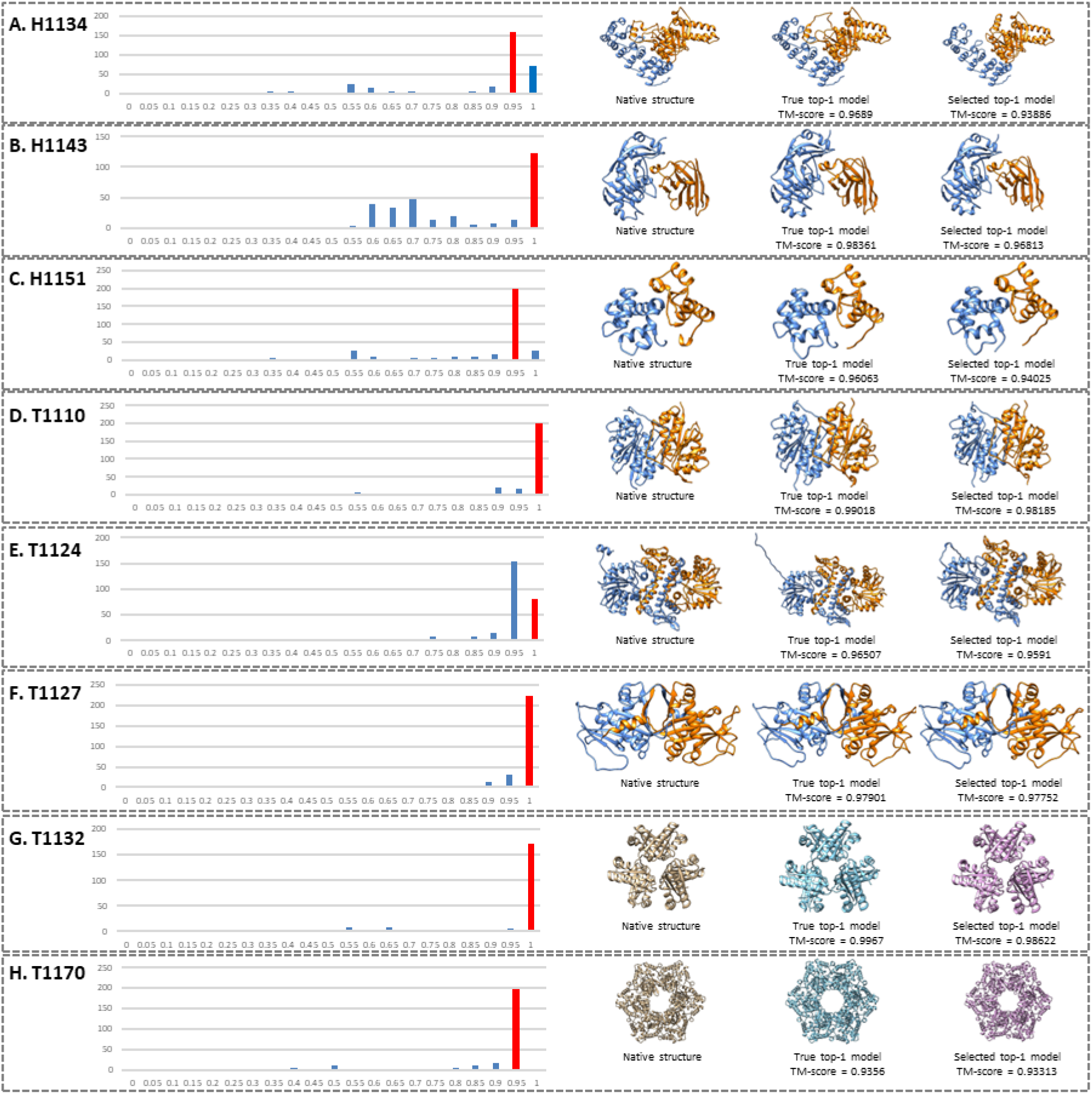
Good examples of applying MULTICOM_qa to rank CASP15 models. Left: the TM-score distribution of the models for each target (the TM-score of the top-1 model selected by MULTICOM_qa comes from the red bin); right: images of the native structure, true top-1 model in the model pool and the top-1 model selected MULTICOM_qa. (**A)** H1134, stoichiometry=A1B1, correlation=0.9877, loss=0.03004; (**B)** H1143, stoichiometry=A1B1, correlation=0.9644, loss=0.01548; (**C)** H1151, stoichiometry=A1B1, correlation=0.9815, loss=0.02038; (**D)** T1110, stoichiometry=A2, correlation=0.8828, loss=0.00833; (**E)** T1124, stoichiometry=A2, correlation=0.9076, loss=0.00597; (**F)** T1127, stoichiometry=A2, correlation=0.7784, loss=0.00149; **G.** T1132, stoichiometry=A6, correlation=0.9323, loss=0.01048; and (**H)** T1170, stoichiometry=A6, correlation=0.9937, loss=0.00247.

**Figure S5.**
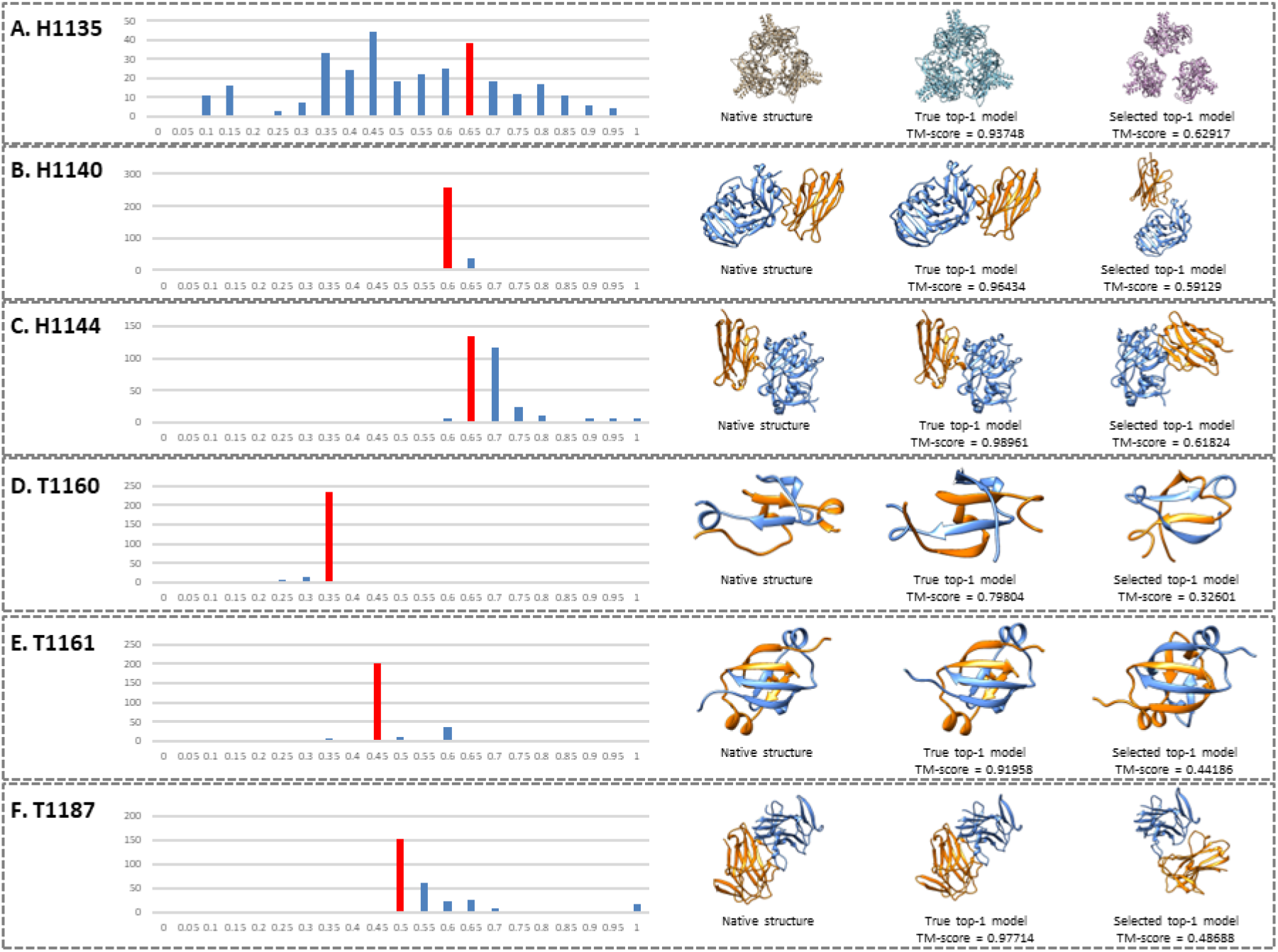
Bad examples of applying MULTICOM_qa to rank CASP15 models. (**A)** H1135, stoichiometry=A9B3, correlation=0.8169, loss=0.30831; (**B)** H1140, stoichiometry=A1B1, correlation=-0.1009, loss=0.37305; (**C)** H1144, stoichiometry=A1B1, correlation=0.1784, loss=0.37137; (**D)** T1160, stoichiometry=A2, correlation=0.129, loss=0.47203; (**E)** T1161, stoichiometry=A2, correlation=-0.1191, loss=0.47772; and (**F)** T1187, stoichiometry=A2, correlation=-0.3537, loss=0.49026.

**Figure S6.**
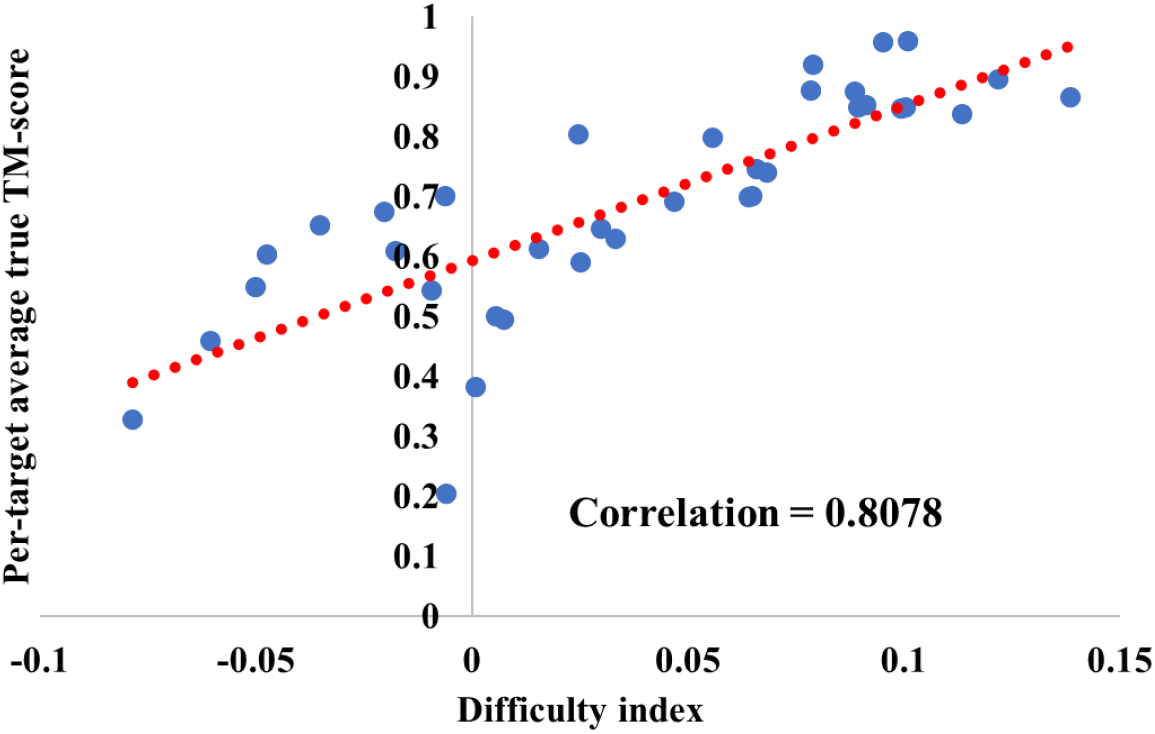
The plot of the average true TM-score of the models of each target against the difficulty index of its models for 36 CASP15 assembly targets. The correlation between the two is 0.8078.

**Figure S7.**
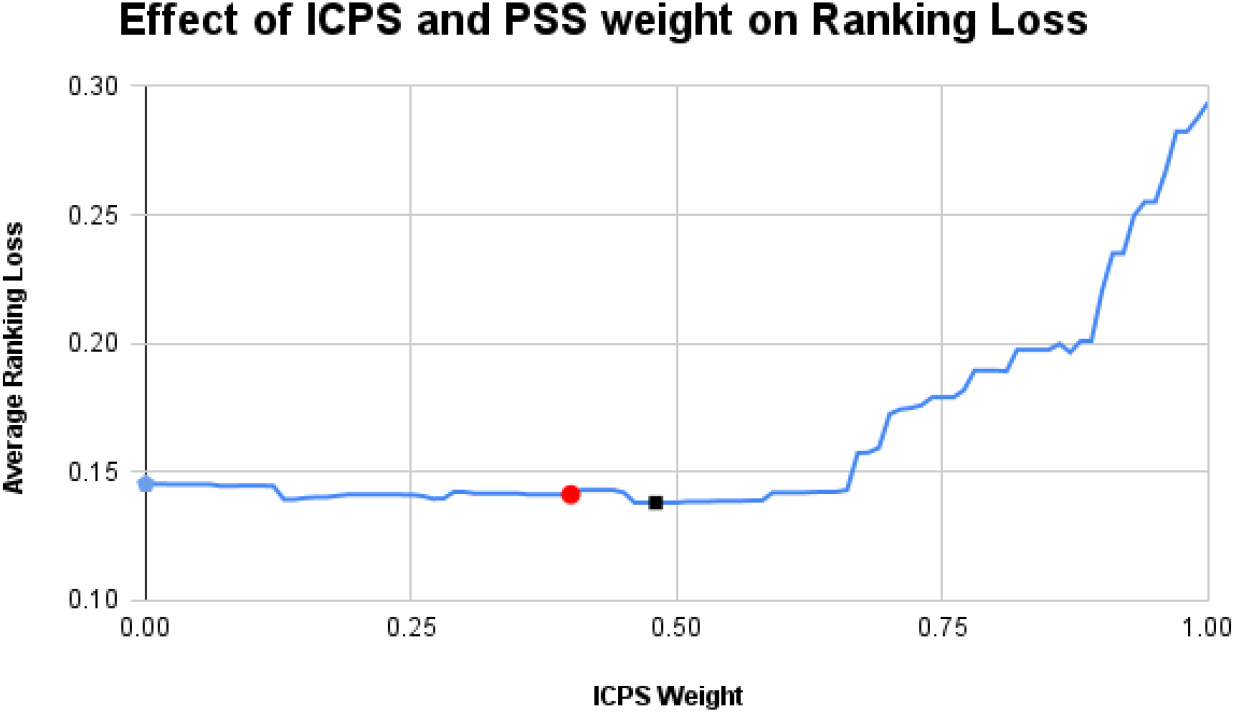
The effect of the weights of ICPS and PSS of MULTICOM_qa on the average per-target ranking loss (PTRL) on the 36 multimer targets of CASP15. The average PTRL is plotted against the weight of ICPS. The weight of PSS is (1 - the weight of ICPS). The weight of ICPS is incremented by 0.01 from 0 to 1. The red point has the loss (0.1413) for the ICPS weight 0.4 used in the CASP15 experiment, the black one is the point with the lowest loss (0.1381) of using ICPS weight 0.48, and the blue one is the point with the loss (0.1454) of using ICPS weight 0 (i.e., only PSS is used).

## Reference

1. Basu, S., & Wallner, B. (2016). DockQ: A Quality Measure for Protein-Protein Docking Models. PLOS ONE, 11(8), e0161879. https://doi.org/10.1371/journal.pone.0161879

2. Bryant, P., Pozzati, G., Zhu, W., Shenoy, A., Kundrotas, P., & Elofsson, A. (2022). Predicting the structure of large protein complexes using AlphaFold and Monte Carlo tree search. Nature Communications, 13(1), Article 1. https://doi.org/10.1038/s41467-022-33729-4

3. Chen, C., Chen, X., Morehead, A., Wu, T., & Cheng, J. (2023). 3D-equivariant graph neural networks for protein model quality assessment. Bioinformatics, 39(1), btad030. https://doi.org/10.1093/bioinformatics/btad030

4. Chen, R., Li, L., & Weng, Z. (2003). ZDOCK: An initial-stage protein-docking algorithm. Proteins: Structure, Function, and Bioinformatics, 52(1), 80–87. https://doi.org/10.1002/prot.10389

5. Chen, X., Morehead, A., Liu, J., & Cheng, J. (2022). DProQ: A Gated-Graph Transformer for Protein Complex Structure Assessment (p. 2022.05.19.492741). bioRxiv. https://doi.org/10.1101/2022.05.19.492741

6. Cheng, J., Choe, M.-H., Elofsson, A., Han, K.-S., Hou, J., Maghrabi, A. H. A., McGuffin, L. J., Menéndez-Hurtado, D., Olechnovič, K., Schwede, T., Studer, G., Uziela, K., Venclovas, Č., & Wallner, B. (2019). Estimation of model accuracy in CASP13. Proteins, 87(12), 1361–1377. https://doi.org/10.1002/prot.25767

7. Cheng, J., Wang, Z., Tegge, A. N., & Eickholt, J. (2009). Prediction of global and local quality of CASP8 models by MULTICOM series. Proteins: Structure, Function, and Bioinformatics, 77(S9), 181–184. https://doi.org/10.1002/prot.22487

8. Dapkūnas, J., Olechnovič, K., & Venclovas, Č. (2021). Modeling of protein complexes in CASP14 with emphasis on the interaction interface prediction. Proteins: Structure, Function, and Bioinformatics, 89(12), 1834–1843. https://doi.org/10.1002/prot.26167

9. Eickholt, J., & Cheng, J. (2012). Predicting protein residue–residue contacts using deep networks and boosting. Bioinformatics, 28(23), 3066–3072. https://doi.org/10.1093/bioinformatics/bts598

10. Evans, R., O’Neill, M., Pritzel, A., Antropova, N., Senior, A., Green, T., Žídek, A., Bates, R., Blackwell, S., Yim, J., Ronneberger, O., Bodenstein, S., Zielinski, M., Bridgland, A., Potapenko, A., Cowie, A., Tunyasuvunakool, K., Jain, R., Clancy, E.,… Hassabis, D. (2022). Protein complex prediction with AlphaFold-Multimer (p. 2021.10.04.463034). bioRxiv. https://doi.org/10.1101/2021.10.04.463034

11. Guo, Z., Liu, J., Skolnick, J., & Cheng, J. (2022). Prediction of inter-chain distance maps of protein complexes with 2D attention-based deep neural networks (p. 2022.06.19.496734). bioRxiv. https://doi.org/10.1101/2022.06.19.496734

12. Hou, J., Wu, T., Cao, R., & Cheng, J. (2019). Protein tertiary structure modeling driven by deep learning and contact distance prediction in CASP13. Proteins, 87(12), 1165–1178. https://doi.org/10.1002/prot.25697

13. Igashov, I., Olechnovič, K., Kadukova, M., Venclovas, Č., & Grudinin, S. (2021). VoroCNN: Deep convolutional neural network built on 3D Voronoi tessellation of protein structures. Bioinformatics, 37(16), 2332–2339. https://doi.org/10.1093/bioinformatics/btab118

14. Jumper, J., Evans, R., Pritzel, A., Green, T., Figurnov, M., Ronneberger, O., Tunyasuvunakool, K., Bates, R., Žídek, A., Potapenko, A., Bridgland, A., Meyer, C., Kohl, S. A. A., Ballard, A. J., Cowie, A., Romera-Paredes, B., Nikolov, S., Jain, R., Adler, J.,… Hassabis, D. (2021). Highly accurate protein structure prediction with AlphaFold. Nature, 596(7873), 583–589. https://doi.org/10.1038/s41586-021-03819-2

15. Olechnovič, K., & Venclovas, Č. (2017). VoroMQA: Assessment of protein structure quality using interatomic contact areas. Proteins: Structure, Function, and Bioinformatics, 85(6), 1131–1145. https://doi.org/10.1002/prot.25278

16. Kandathil, S. M., Greener, J. G., & Jones, D. T. (2019). Prediction of interresidue contacts with DeepMetaPSICOV in CASP13. Proteins: Structure, Function, and Bioinformatics, 87(12), 1092–1099. https://doi.org/10.1002/prot.25779

17. Kryshtafovych, A., Monastyrskyy, B., & Fidelis, K. (2014). CASP prediction center infrastructure and evaluation measures in CASP10 and CASP ROLL. Proteins, 82 Suppl 2, 7–13. https://doi.org/10.1002/prot.24399

18. Kryshtafovych, A., Schwede, T., Topf, M., Fidelis, K., & Moult, J. (2019). Critical assessment of methods of protein structure prediction (CASP)—Round XIII. Proteins: Structure, Function, and Bioinformatics, 87(12), 1011–1020. https://doi.org/10.1002/prot.25823

19. Kuhn, H. W. (1955). The Hungarian method for the assignment problem. Naval Research Logistics Quarterly, 2(1-2), 83–97. https://doi.org/10.1002/nav.3800020109

20. Kwon, S., Won, J., Kryshtafovych, A., & Seok, C. (2021). Assessment of protein model structure accuracy estimation in CASP14: Old and new challenges. Proteins: Structure, Function, and Bioinformatics, n/a(n/a). https://doi.org/10.1002/prot.26192

21. Larsson, P., Skwark, M. J., Wallner, B., & Elofsson, A. (2009). Assessment of global and local model quality in CASP8 using Pcons and ProQ. Proteins, 77 Suppl 9, 167–172. https://doi.org/10.1002/prot.22476

22. Lensink, M. F., Brysbaert, G., Mauri, T., Nadzirin, N., Velankar, S., Chaleil, R. A. G., Clarence, T., Bates, P. A., Kong, R., Liu, B., Yang, G., Liu, M., Shi, H., Lu, X., Chang, S., Roy, R. S., Quadir, F., Liu, J., Cheng, J.,… Wodak, S. J. (2021). Prediction of protein assemblies, the next frontier: The CASP14-CAPRI experiment. Proteins: Structure, Function, and Bioinformatics, n/a(n/a). https://doi.org/10.1002/prot.26222

23. Lensink, M. F., Velankar, S., Baek, M., Heo, L., Seok, C., & Wodak, S. J. (2018). The challenge of modeling protein assemblies: The CASP12-CAPRI experiment. Proteins: Structure, Function, and Bioinformatics, 86(S1), 257–273. https://doi.org/10.1002/prot.25419

24. Lensink, M. F., Velankar, S., Kryshtafovych, A., Huang, S.-Y., Schneidman-Duhovny, D., Sali, A., Segura, J., Fernandez-Fuentes, N., Viswanath, S., Elber, R., Grudinin, S., Popov, P., Neveu, E., Lee, H., Baek, M., Park, S., Heo, L., Lee, G. R., Seok, C.,… Wodak, S. J. (2016). Prediction of homoprotein and heteroprotein complexes by protein docking and template-based modeling: A CASP-CAPRI experiment. Proteins: Structure, Function, and Bioinformatics, 84(S1), 323–348. https://doi.org/10.1002/prot.25007

25. Liu, J., Wu, T., Guo, Z., Hou, J., & Cheng, J. (2022). Improving protein tertiary structure prediction by deep learning and distance prediction in CASP14. Proteins: Structure, Function, and Bioinformatics, 90(1), 58–72. https://doi.org/10.1002/prot.26186

26. Liu, J., Zhao, K., & Zhang, G. (2023). Improved model quality assessment using sequence and structural information by enhanced deep neural networks. Briefings in Bioinformatics, 24(1), bbac507. https://doi.org/10.1093/bib/bbac507

27. Lu, H., Lu, L., & Skolnick, J. (2003). Development of Unified Statistical Potentials Describing Protein-Protein Interactions. Biophysical Journal, 84(3), 1895–1901.

28. McGuffin, L. J., Aldowsari, F. M. F., Alharbi, S. M. A., & Adiyaman, R. (2021). ModFOLD8: Accurate global and local quality estimates for 3D protein models. Nucleic Acids Research, 49(W1), W425–W430. https://doi.org/10.1093/nar/gkab321

29. Moult, J., Fidelis, K., Kryshtafovych, A., Schwede, T., & Tramontano, A. (2016). Critical assessment of methods of protein structure prediction: Progress and new directions in round XI. Proteins: Structure, Function, and Bioinformatics, 84(S1), 4–14. https://doi.org/10.1002/prot.25064

30. Moult, J., Pedersen, J. T., Judson, R., & Fidelis, K. (1995). A large-scale experiment to assess protein structure prediction methods. Proteins: Structure, Function, and Bioinformatics, 23(3), ii–iv. https://doi.org/10.1002/prot.340230303

31. Mukherjee, S., & Zhang, Y. (2009). MM-align: A quick algorithm for aligning multiple-chain protein complex structures using iterative dynamic programming. Nucleic Acids Research, 37(11), e83. https://doi.org/10.1093/nar/gkp318

32. Pierce, B., & Weng, Z. (2007). ZRANK: Reranking protein docking predictions with an optimized energy function. Proteins, 67(4), 1078–1086. https://doi.org/10.1002/prot.21373

33. Pierce, B., & Weng, Z. (2008). A combination of rescoring and refinement significantly improves protein docking performance. Proteins, 72(1), 270–279. https://doi.org/10.1002/prot.21920

34. Quadir, F., Roy, R. S., Soltanikazemi, E., & Cheng, J. (2021). DeepComplex: A Web Server of Predicting Protein Complex Structures by Deep Learning Inter-chain Contact Prediction and Distance-Based Modelling. Frontiers in Molecular Biosciences, 8, 827. https://doi.org/10.3389/fmolb.2021.716973

35. Roy, R. S., Quadir, F., Soltanikazemi, E., & Cheng, J. (2022). A deep dilated convolutional residual network for predicting interchain contacts of protein homodimers. Bioinformatics, 38(7), 1904–1910. https://doi.org/10.1093/bioinformatics/btac063

36. Senior, A. W., Evans, R., Jumper, J., Kirkpatrick, J., Sifre, L., Green, T., Qin, C., Žídek, A., Nelson, A. W. R., Bridgland, A., Penedones, H., Petersen, S., Simonyan, K., Crossan, S., Kohli, P., Jones, D. T., Silver, D., Kavukcuoglu, K., & Hassabis, D. (2019). Protein structure prediction using multiple deep neural networks in the 13th Critical Assessment of Protein Structure Prediction (CASP13). Proteins: Structure, Function, and Bioinformatics, 87(12), 1141–1148. https://doi.org/10.1002/prot.25834

37. Senior, A. W., Evans, R., Jumper, J., Kirkpatrick, J., Sifre, L., Green, T., Qin, C., Žídek, A., Nelson, A. W. R., Bridgland, A., Penedones, H., Petersen, S., Simonyan, K., Crossan, S., Kohli, P., Jones, D. T., Silver, D., Kavukcuoglu, K., & Hassabis, D. (2020). Improved protein structure prediction using potentials from deep learning. Nature, 577(7792), 706–710. https://doi.org/10.1038/s41586-019-1923-7

38. Studer, G., Tauriello, G., & Schwede, T. (2022). CASP15 EMA. https://predictioncenter.org/casp15/doc/presentations/Day4/Assessment_EMA_GStuder.pdf

39. Wallner, B., & Elofsson, A. (2006). Identification of correct regions in protein models using structural, alignment, and consensus information. Protein Science: A Publication of the Protein Society, 15(4), 900. https://doi.org/10.1110/ps.051799606

40. Wang, S., Sun, S., Li, Z., Zhang, R., & Xu, J. (2017). Accurate De Novo Prediction of Protein Contact Map by Ultra-Deep Learning Model. PLOS Computational Biology, 13(1), e1005324. https://doi.org/10.1371/journal.pcbi.1005324

41. Wang, X., Flannery, S. T., & Kihara, D. (2021). Protein Docking Model Evaluation by Graph Neural Networks. Frontiers in Molecular Biosciences, 8. https://www.frontiersin.org/articles/10.3389/fmolb.2021.647915

42. Wang, X., Terashi, G., Christoffer, C. W., Zhu, M., & Kihara, D. (2020). Protein docking model evaluation by 3D deep convolutional neural networks. Bioinformatics (Oxford, England), 36(7), 2113–2118. https://doi.org/10.1093/bioinformatics/btz870

43. Wang, Z., Eickholt, J., & Cheng, J. (2011). APOLLO: A quality assessment service for single and multiple protein models. Bioinformatics, 27(12), 1715–1716. https://doi.org/10.1093/bioinformatics/btr268

44. Xie, Z., & Xu, J. (2021). Deep graph learning of inter-protein contacts. Bioinformatics (Oxford, England), btab761. https://doi.org/10.1093/bioinformatics/btab761

45. Yan, Y., & Huang, S.-Y. (2021). Accurate prediction of inter-protein residue–residue contacts for homo-oligomeric protein complexes. Briefings in Bioinformatics, bbab038. https://doi.org/10.1093/bib/bbab038

46. Yang, J., Anishchenko, I., Park, H., Peng, Z., Ovchinnikov, S., & Baker, D. (2020). Improved protein structure prediction using predicted interresidue orientations. Proceedings of the National Academy of Sciences, 117(3), 1496–1503. https://doi.org/10.1073/pnas.1914677117

47. Zemla, A. (2003). LGA: A method for finding 3D similarities in protein structures. Nucleic Acids Research, 31(13), 3370. https://doi.org/10.1093/nar/gkg571

48. Zhang, Y., & Skolnick, J. (2004). SPICKER: A clustering approach to identify near-native protein folds. Journal of Computational Chemistry, 25(6), 865–871. https://doi.org/10.1002/jcc.20011

49. Zhang, Y., & Skolnick, J. (2004, October 8). Scoring function for automated assessment of protein structure template quality—Zhang—2004—Proteins: Structure, Function, and Bioinformatics—Wiley Online Library. https://onlinelibrary.wiley.com/doi/10.1002/prot.20264

50. Zheng, W., Li, Y., Zhang, C., Pearce, R., Mortuza, S. M., & Zhang, Y. (2019). Deep-learning contact-map guided protein structure prediction in CASP13. Proteins, 87(12), 1149–1164. https://doi.org/10.1002/prot.25792

